# Visuomotor coordination on the road: low-dimensional representations reveal adaptive, context-dependent reductions in the dimensionality of natural driving behavior

**DOI:** 10.64898/2026.02.28.707665

**Authors:** John Madrid-Carvajal, Shadi Derakhshan, Peter König

## Abstract

The brain’s ability to transform complex, high-dimensional sensory and motor inputs into coordinated, goal-directed behavior remains a central challenge in neuroscience research. Current research suggests that behavior is generated through patterns within a low-dimensional structure. What visuomotor patterns underlie complex, unconstrained behaviors such as driving, and how stereotypical they are, remain poorly understood. Driving involves complex perception-action interactions, which we hypothesize are specific to the driver’s role and adapt to task demands. Here, we unfold these dynamics using an immersive virtual drive that recreates ten hazardous on-road events. We collected eye, head, and vehicle movements from participants assigned to either a manual or an autonomous driving condition. From the behavioral movements of 284 drivers, we constructed a common time-resolved behavioral state space based on collective variance, computed via principal component analysis across participants. The obtained low-dimensional manifold thus represents generalized visuomotor coordination strategies. We then characterized the evolving low-dimensional structure with cosine similarity and various entropy-based measures of effective dimensionality. We tested whether early components contained condition-specific information using discriminant analysis. Across the entire drive, a few components accounted for most of the variance in behavior, and effective dimensionality decreased reliably around critical events, indicating tighter coordination between perceptual and motor variables. During periods of low effective dimensionality, the contributions of eye, head, steering, and vehicle heading reorganized toward early dimensions in a task-dependent manner. As a result, behaviors became more distinct across driving conditions, enabling better classification of driving modes using only the first two components. These results show that naturalistic driving is supported by shared low-dimensional visuomotor coordination strategies that are flexibly reshaped by contextual demands, and provide a principled behavioral framework for linking neural manifolds, human driver models, and the design of adaptive autonomous vehicles.

**Author summary:** Human behavior is complex. Driving, for instance, means constantly turning what we see into the right movements of our head, hands, and feet. Research using constrained repeated actions suggests that these seemingly complex behaviors are highly coordinated and can be described by a relatively small number of variables. We want to know whether these aspects also hold during natural, unconstrained tasks. We asked people to complete a virtual drive comprising ten sudden hazards, such as animals and pedestrians on the road. Some participants drove the car, while others sat in a self-driving car. We recorded where people looked, how they moved their heads, and steered the vehicle. Then, we used statistical methods to reduce the data’s dimensionality and looked for patterns that could explain all these signals together. We found that a small number of patterns were enough to describe behavior. These patterns became even simpler and more linked around hazards, differentiating autonomous from manual driving behavior. Our results suggest that even complex driving behavior follows a few basic coordination strategies that become more pronounced around specific situations. These basic dimensions contain important details that are unique to certain groups, ultimately helping us better understand and compare human behavior.

## Introduction

A major challenge when studying visuomotor coordination in large natural settings concerns the inherent complexity of actions. Ordinary everyday actions involve remarkably complex behaviors implicating many degrees of freedom (DoF). The DoF problem concerns the challenge of effectively controlling a system with so many variables that can move independently during natural behavior [1–3]. When walking, for instance, the central nervous system exerts control over many DoF within the musculoskeletal system, coordinating movement across multiple levels [4]. From the primary effectors (i.e., the muscles) to the resulting coordinated motion of the limbs, trunk, arms, head, and eyes, to the underlying bodily processes that occur along with them. Despite considerable progress in understanding how the central nervous system coordinates this high-dimensional movement system, the specific strategies that it utilizes to effectively control and solve the DoF problem during natural behavior remain an open question.

A second aspect that makes behavior complex concerns its high variability. Variability is often attributed to the fact that every single movement can be carried out in multiple ways, where even slight variations in body posture can lead to different outcomes [5, 6]. It arises during motor preparation [7] and is likely linked to errors in the sensory estimates of the external world [8]. This variability has typically been studied separately within subjects and compared across subjects to assess similarities in the underlying mechanisms (i.e., stereotypy) [9–15]. Quantifying behavioral stereotypy (whether the central nervous system utilizes consistent strategies to organize behavior despite its extensive redundancy) requires developing solid frameworks that relate the multiscale and distributed dynamics inherent to behavior [16, 17]. Equally important are methods to characterize the structure of behavior itself [18–23]. Notably, these methods have been predominantly applied to animal behavior. A key concept in this context is dimensionality, which quantifies the size of the subspace effectively used for behavior relative to the space of all possible movements [16, 19, 24]. Essentially, higher variability in behavior may reflect a higher dimensionality, indicating a broader range of directions in the movement space. Conversely, lower variability suggests movement is restricted to a narrower set of control strategies. Thus, understanding how the central nervous system produces coordinated behavior despite redundancy implies determining its dimensionality.

The difficulty of analyzing complex, variable behavior has led contemporary research to utilize dimensionality reduction techniques to reduce high-dimensional behavioral and neural data into a low-dimensional structure. Common methods include linear and nonlinear variants, with linear approaches such as principal component analysis (PCA) proving useful [25–30]. PCA reduces data complexity by providing a set of eigenvectors that describes as much variability as possible with fewer components, capturing key patterns of covariation in the original data. This process yields two primary outputs: a set of orthogonal components (*eigenvectors*) and their associated *eigenvalues*, which quantify the magnitude of the variance each component captures [31–33]. The resulting principal components (PCs) are mutually independent linear combinations of the original variables that maximize the total variance explained. Although PCA reduces data complexity, quantifying the effective dimensionality (ED) is not trivial. One way to address this issue is to utilize the full eigenvalue spectrum, whose magnitudes indicate the amount of variance retained by each component. We then measure ED using, for instance, entropy-based estimators which yield an interpretable, information-driven measure of the equivalent number of independent dimensions required to account for the overall pattern of covariation in the data [24, 34, 35] (see Methods section). Consequently, a low ED value indicates tighter cross-variable coordination within the system.

In neuroscience, PCA has been used to directly relate behavioral data to its underlying neural activity, shedding light on the type of transformations involved in the production and control of movement [36–39]. The core hypothesis suggests that, despite its many DoF and variability, behavior generation is constrained to a low-dimensional subspace or manifold that captures the essential information for the planning, execution, and learning of movements [28, 40, 41]. These manifolds have successfully explained the organization of diverse behaviors, including locomotion [42–44] and reaching/grasping [25, 27, 45–47]. Moreover, this low-dimensional structural organization has been consistently found in non-human primates [48, 49], mice [29, 50, 51], Caenorhabditis elegans [22], zebrafish [52], and birds [53]. A predominant aspect of the literature, however, is its reliance on stereotyped tasks and its focus primarily on non-human species. Furthermore, the application of these methods to quantify complex natural human action is sparse. For instance, the dimensionality of visuomotor coordination involving eye, hand, and head kinematics is understudied. A recent study investigating naturalistic behavior revealed a synergistic coupling of the eye-head and eye-hand systems during a pick-up and drop-off task [54]. Thus, applying dimensionality reduction to understand human visuomotor coordination in natural environments highlights a frontier for the study of human behavior.

Driving is a complex task that provides a rich context for investigating visuomotor coordination. It is a skill involving perceptually guided locomotion by means of a vehicle. Driving is initiated and motivated by the goal of reaching a specific destination and maintained by concurrent behaviors where *“the seeing and the doing are merged in the same experience”* [55]. Understanding driver behavior entails discerning the set of behavioral schemas that underlie this complex skill, examining how sensory and motor features interact, and identifying the low-dimensional transformations that generate behavior. The inherent unpredictability of real-world driving poses a significant challenge for visuomotor systems, which must adapt rapidly to maintain safe and effective control. In this context, coordinated, skillful driving involves overcoming superfluous DoF by translating them into goal-oriented behaviors. A transformation of high-dimensional behavioral variables into common and synchronized features underlying the production of skilled behavior [56, 57]. A key question that arises is how many DoF are truly required and which dimensions hold the relevant information to facilitate driving.

Here, we test the hypothesis that the rich, high-dimensional visuomotor coordination supporting naturalistic driving lies on a low-dimensional manifold. Specifically, we investigate whether eye-head-vehicle coordination reveals stereotyped patterns that are shared across drivers yet distinct between driving modes. To address this, we analyzed eye-tracking and behavioral data recorded during a simulated virtual drive comparing manual control versus autonomous operation, featuring ten safety-relevant “critical events” [58]. Crucially, we focused on variability across participants to investigate inter-individual differences in coordination strategies rather than mere movement magnitudes. By applying dimensionality reduction to the behavioral variables, we traced how these coordination patterns evolve over time and identified the variables contributing most to the dominant modes of variation. Finally, we employed discriminant analysis to quantify whether the resulting low-dimensional representation retains sufficient information to reliably distinguish between driving conditions and characterize behavior around critical events. The results reveal a low-dimensional organization of visuomotor coordination that differs meaningfully between driving conditions.

## Materials and methods

### Research design

To understand the low-dimensional dynamics underlying complex natural behavior, we conducted a driving simulation using virtual reality (VR) (Fig 1A). During the drive, participants encountered ten critical hazardous events involving a sudden appearance of predefined driving hazards, such as a reindeer dashing through the street (Fig 1B and S1 Video). As in other real-world critical driving scenarios, the simulation included hazards such as animals, pedestrians, vehicles, and landslides that posed driving challenges. The experiment consisted of two conditions: manual or autonomous driving (Fig 1C). While the autonomous condition required participants to sit and experience a fully autonomous drive (hands-off wheel), the manual drive gave participants complete control of the car’s steering wheel and pedals, and they were instructed to perform the drive as they would in real life (hands-on wheel). In both conditions, participants were seated behind the wheel. By replicating real-world driving scenarios using VR, we could safely collect and compare the dynamics of complex driving behavior across different conditions.

**Fig 1.**
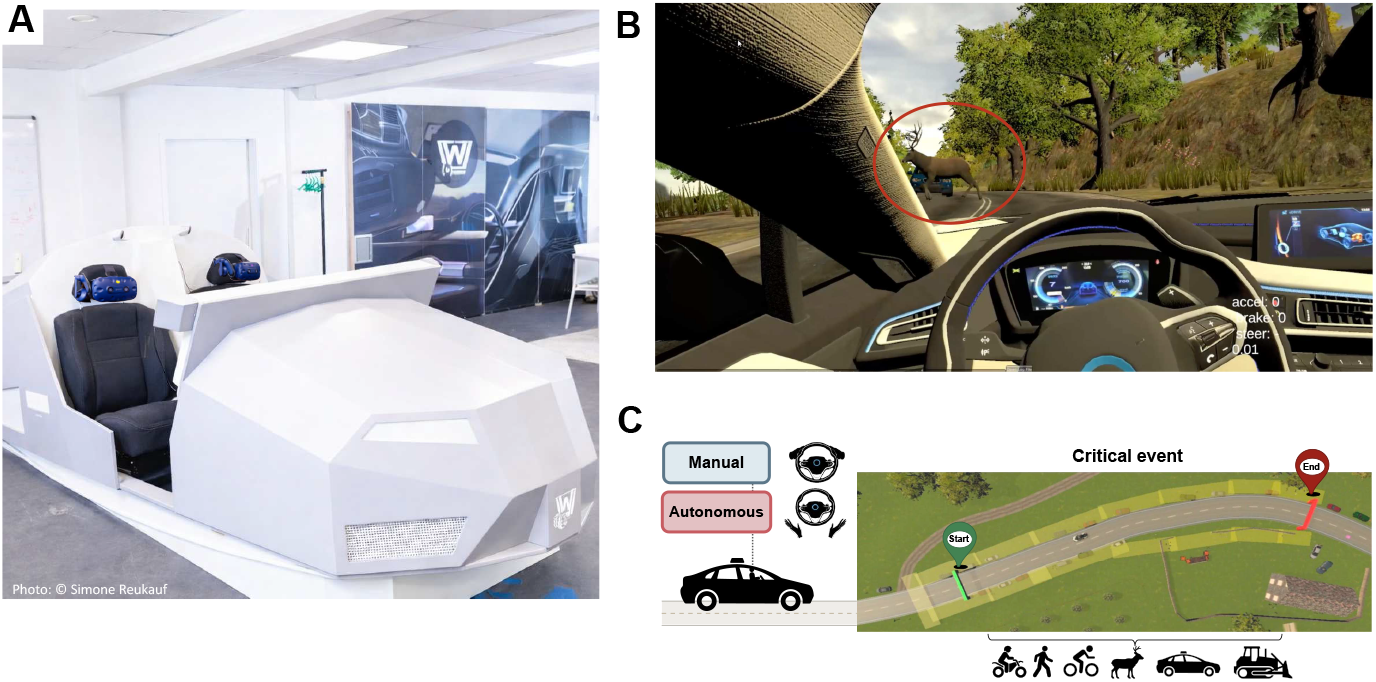
Experimental setup and virtual environment. (A) The wooden car simulator where participants sat wearing the Head-Mounted Display (HMD). (B) Participants’ immersive view in virtual reality (VR), showing a reindeer dashing through the street—the red circle was not visible to participants. (C) An aerial representation of the manual hands-on wheel (blue) and autonomous hands-off wheel (red) driving conditions in VR during a critical event from start to end.

We collected various behavioral metrics during the driving simulation to analyze participants’ responses and compare behavioral performance under manual versus autonomous driving. Specifically, we collected the trajectories of the eye, head, and car in 3D space, along with their positions. Additionally, the car input data included steering, acceleration, and braking movements. These metrics, which characterize the perceptual and action schemas employed in driving, enabled us to examine the dimensionality of complex human behavior.

### Materials and procedure

In this study, we analyzed eye-tracking and behavioral data collected during a simulated virtual drive. To provide a realistic experience, the experiment integrated a Head-Mounted Display (HMD) for stimulus presentation and a wooden car. The data was collected using an HTC Vive Pro Eye with integrated Tobii eye tracking, 110^◦^ field of view, 1440 × 1600 pixel resolution per eye, and 0.5–1.1^◦^ accuracy (https://developer.vive.com/resources/hardware-guides/vive-pro-eye-specs-user-guide/). The integrated eye-tracking system used the SDK SRanipal v1.1.0.1 (https://developer.vive.com/resources/vive-sense/eye-and-facial-tracking-sdk). The VR scene maintained an average frame rate of 88 FPS and was created in Unity 3D (version 2019.3.0f3, 64 bit; https://unity.com/). The driving scene was adapted from a publicly available VR toolkit created to investigate autonomous driving [59]. The virtual driving environment was designed to confront drivers with a real-life driving experience; it covers approximately 11 km^2^ and features multiple animated pedestrians, cars, and animals, as well as ten critical hazardous situations (for details, see [58]). Steering and braking inputs were recorded using a Fanatec CSL Elite Racing Wheel (PS4™) with an adjustable turning angle of 90-1080^◦^, 970 g metal rim, 300mm realistic diameter, and genuine leather grips and CSL Elite Pedals with 12-bit resolution potentiometers and two pedal sockets (https://fanatec.com/eu-en/csl-elite-pedals?number=CSL_EP).

### Ethics statement

The study was conducted in accordance with the Declaration of Helsinki and approved by the Institutional Ethics Committee of the University of Osnabrück (approval number: Ethik37-2021), and written informed consent was obtained from all participants involved in this study before participation.

### Participants

This study involved 2,840 visitors during a year-long exhibition at the Deutsches Museum in Bonn (Germany). Participants could withdraw at any time without having to provide any explanation. As a result, many datasets were incomplete because participants withdrew from the study at various stages. For this reason, the present analysis included 284 participants who conducted either a manual (*N* = 131) or an autonomous (*N* = 153) virtual drive and had valid eye-tracking data. Demographic information was available only for 168 of the participants included (125 male, 37 female, 6 intersex) who completed the follow-up questionnaire. The mean age of this subset of participants was 23.4 years (*SD* = 15.6). No demographics-based analysis was performed in this study.

### Segmentation and principal components analysis

We investigated the low-dimensional behavioral dynamics underlying driving using a structured methodology for data segmentation and dimensionality reduction via principal component analysis (PCA) [31–33]. The analysis was conducted in Python, and the results were compared with those from other statistical software [60]. Initially, the dataset comprising several behavioral variables, including eye, head, car, and steering behavior over time, was organized into a three-dimensional block structure, where each dimension represented a behavioral variable at a timepoint for each participant (Fig 2A). From this structure, we first segmented the high-dimensional dataset into time segments for the complete duration of the ride, yielding a subjects × features matrix (Fig 2B). For each time segment, PCA was applied to transform the high-dimensional parameter space (i.e., the multidimensional set of features characterizing the dataset) into a simplified and interpretable representation (Fig 2C). Before PCA, we standardized the data to have all variables in a similar unit range, treating each variable equally during analysis regardless of its original scale [33, 61]. To characterize how the course of dimensionality evolved within a single event, we quantified variable-dimension associations as Pearson correlations between each variable and the PC scores at each timepoint. For analysis, we focused on strategic timepoints such as the event onset and offset, plus two additional timepoints that were identified based on the variance explained by the first two components: the peak with the highest variance after onset and the decline with the lowest variance after onset. Two-sided *p*-values were obtained from the corresponding *t*-statistic. This process yielded an interpretable low-dimensional representation of the inherent behavioral dynamics that captured the main patterns of covariation in the data.

**Fig 2.**
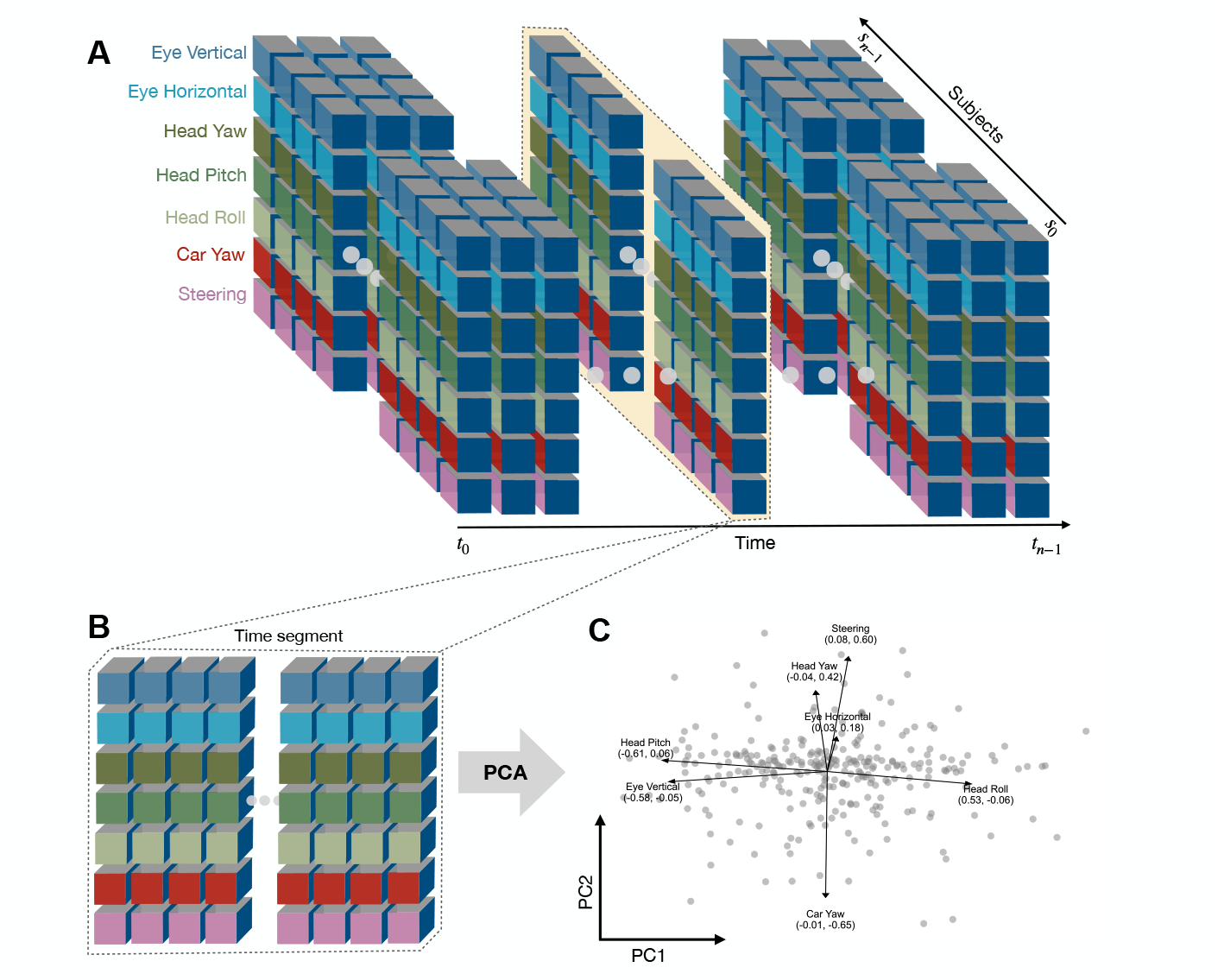
High-dimensional data segmentation and dimensionality reduction procedure. (A) An illustration of the multidimensional dataset composed of several behavioral parameters over time (x-axis), including eye, head, car, and steering movements, and color-coded for clarity. Each vertical block (y-axis) contains all behavioral data points for each subject (gray z-axis) within a specific timepoint from *t*_0_ to *t*_*n*−1_. (B) A highlighted timepoint showing the extracted segment to which principal component analysis (PCA) is applied. (C) Resulting biplot after PCA, showing the distribution of the data after projecting it into the first two PCs (PC1 and PC2). Each gray circle in the biplot represents a subject, and the vectors indicate the contribution and correlation of each original variable in the reduced dimensional space; the larger the vector, the more it contributed to the corresponding PC.

### Data preprocessing

The behavioral variables recorded in Unity consisted of the x-, y-, and z-axis unit vectors for the position and direction of eye movements (Fig 3). These 3D vectors encode rotation angles (in degrees) along each axis. Head and car rotations were measured as quaternions, that is, four-tuples (x, y, z, w) that encode the spatial orientation and rotation in 3D space [62]. To measure participants’ steering actions, steering input was collected in radians from the game-ready Fanatec steering wheel device. Eye unit vectors (position and direction) were collected in the local head reference frame, while head and car rotations were measured in the global reference frame. We therefore obtained local head movements by subtracting head motion from car motion. In an additional step, we aligned each participant’s data to a standard reference time based on the car’s position from a predetermined autonomous drive and resampled it to 50 Hz. This process ensured that events were temporally matched across participants and had similar durations. The following subsections detail how gaze, head, and car signals were converted to angles.

**Fig 3.**
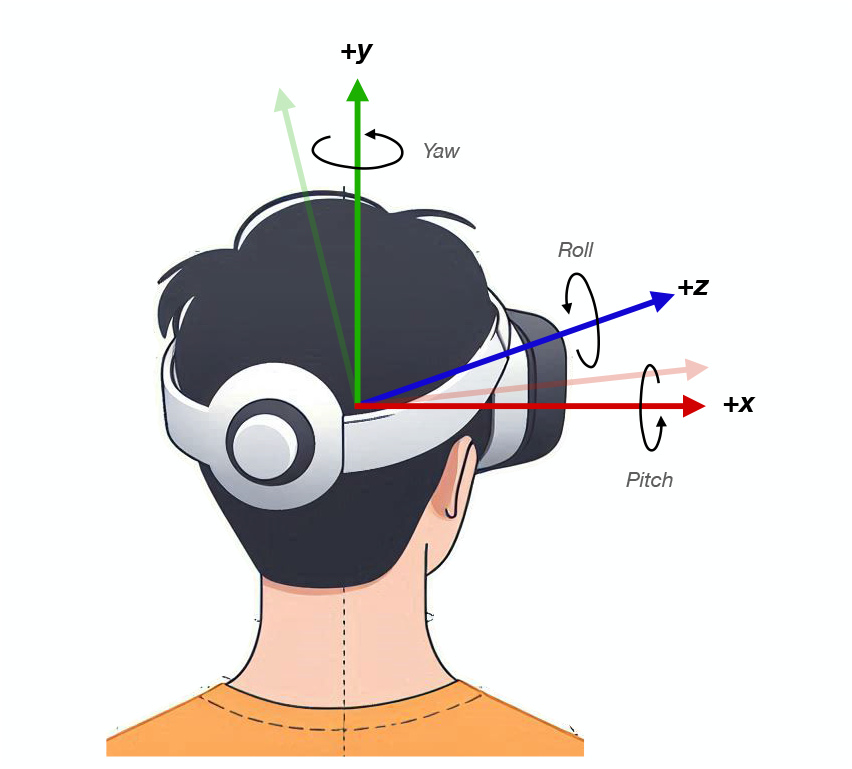
Unity 3D left-handed coordinate system representing rotations and orientations in 3D space. The z-axis points forward (blue), the x-axis points towards the right (red), and the y-axis points upwards (green). The dimmed red and green arrows depict Unity’s counterclockwise direction of rotation along +z from +x to +y. Circular arrows illustrate the Euler angles for head movements: roll (side-to-side tilt), pitch (up-and-down nod), and yaw (left-to-right turn).

To express all measurements in a common unit, we converted the preprocessed signals to degrees using Euler angles (see the following subsections). We then used a robust Mahalanobis-distance-based procedure for multivariate outlier detection to identify extreme outliers in the data. Because eye, head, and car movements differ in scale and type, outliers were identified and handled within each category by applying this procedure separately to each movement category (horizontal and vertical eye, head pitch, yaw, and roll, etc.).

### Gaze data

Unity uses a left-handed coordinate system with the y-axis pointing upward, unlike the right-handed system used in most 3D modeling environments. Rotations follow the “z-x-y” order. Together, the upward y-axis and this rotation order determine how rotations map to Euler angles [63, 64]; these conventions were applied when converting Unity output to angles. We converted the 3D unit vectors of gaze direction (x, y, z) into Euler angles to determine horizontal and vertical eye movements in degrees at each timepoint using Eq. (1) and Eq. (2), respectively. Horizontal angle used the arctangent of *x/z*; vertical angle used the arctangent of *y/z*. In these expressions, x is the lateral component, y the vertical component, and z the depth component of the unit vector.

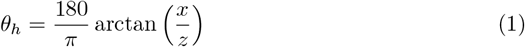

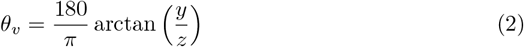

### Head data

Head data was obtained from the Head Mounted Display (HMD) participants wore during the experiment. We converted the head quaternions to Euler angles using the `Rotation` class from the `scipy.spatial.transform` package. This process involved two steps: we first transformed global head quaternions into local head quaternions, then converted the local quaternions to Euler angles. For the first step, we computed the inverse of the parent (car) quaternions. With the global head quaternion 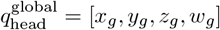 and the parent quaternion *q*_car_ = [*x*_*p*_, *y*_*p*_, *z*_*p*_, *w*_*p*_], the inverse of the unit parent quaternion is given by Eq. (3). The local head quaternion 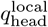 is then obtained using Eq. (4) as the product of the global head quaternion and the parent inverse.

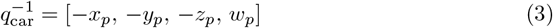

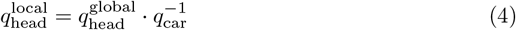

We then converted the local head quaternions to Euler angles using the `as euler` property of the `Rotation` class, with the sequence matched to Unity’s z-x-y order as in Eq. (5). The resulting angles correspond to roll (*ϕ*), pitch (*θ*), and yaw (*ψ*), in that order, where roll represents a rotation along the z-axis, pitch along the x-axis, and yaw along the y-axis. For completeness, the explicit conversion formulas are given in Eqs. (6)–(8). Finally, we converted these angles from radians to degrees using Eq. (9).

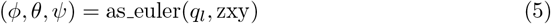

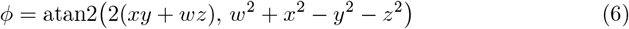

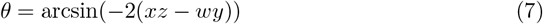

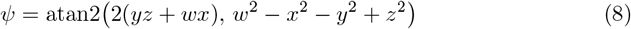

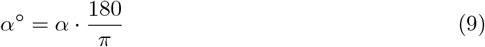

### Car data

The car positions and rotations were obtained as quaternions from the car’s motion in the virtual environment. As for gaze and head data, we used Eqs. (5)–(9) to convert the car rotation quaternions to Euler angles in degrees. Steering was recorded in radians and converted to degrees using Eq. (9).

### Statistical analysis

We performed a detailed analysis of the PCA results to investigate shifts in variance over the duration of the experiment. For each principal component (PC), explained variance was aligned to event onset across all events (*N* = 10). We then compared the before-onset (BO) and after-onset (AO) windows using a ± 5-second window around onset (excluding *t* = 0). We tested whether PC variance in the BO window differed from that in the AO window across all event × PC pairs using an inverse-variance-weighted chi-square test (null hypothesis: BO = AO variance). We expected the early dimensions to explain more variance after onset. Consequently, the initial dimensions would account for a greater share of the variance, and the variance in later dimensions would decrease. A similar analysis was performed to investigate the differences in effective dimensionality (ED) around onset. We computed the BO and AO means (and standard errors) of ED (null hypothesis: BO = AO dimensionality). For both variance and ED, we report the weighted mean change and its 95% confidence interval (CI). These analyses quantify event-evoked shifts in variance and ED and assess the consistency of the underlying dynamics across events.

### Cosine similarity and effective dimensionality

The quality of representation of each variable within each principal component (PC) was assessed using the squared cosine 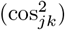 defined in Eq. (10), which quantifies the squared correlation between a variable and a PC.

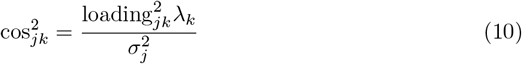

Where loading_*jk*_ is the loading of variable *j* on component *k, λ*_*k*_ is the eigenvalue of component *k*, and *σ*_*j*_ is the standard deviation of variable *j*.

We obtained effective dimensionality (ED) from the eigenvalue spectrum at each timepoint using several entropy-based estimators [34]. In our analysis, we employed all four estimators: *n*_1_, *n*_2_, *n*_*C*_, and *n*_∞_. Each captures different characteristics of the eigenspectrum. *n*_1_ provides a balanced assessment, considering the distribution of variance relatively evenly across dimensions. *n*_2_ is more sensitive to the dominant eigenvalues, emphasizing the contribution of dimensions with the highest variance while discounting smaller ones. *n*_*C*_ is a variance-based estimator that quantifies the dispersion of the eigenvalues, defining dimensionality as the total number of variables minus a penalty term that increases as the distribution of variance becomes more unequal. Finally, *n*_∞_ denotes the limiting case in which only the largest eigenvalue is considered, providing insight into dimensionality dominated by a single factor. ED thus provides an interpretable measure of the number of independent, orthogonal dimensions required to account for the overall covariation pattern observed in the data; a lower ED indicates tighter cross-variable coordination.

### Discriminant analysis and cross-validated AUC

After applying PCA and reducing the behavioral variables to the first two principal components, we tested whether this compact, low-dimensional representation of multi-signal behavior captures reliable condition separation, both geometrically and predictively. To ensure comparability of axes across time, the 2D loadings matrix was aligned to a reference basis using orthogonal Procrustes rotation with reflection suppression, followed by component-wise sign continuity. Condition separation was quantified directly in this aligned PC plane by computing the class centroid for Autonomous (label 0) and Manual (label 1), forming the unit discriminant axis along the centroid difference, and projecting individual PC scores onto this axis. Group-mean projections with bootstrap 95% CI were obtained to report the discriminant score. The Euclidean distance between class centroids yields the reported centroid-distance (with bootstrap CI). For classification performance, we trained a logistic regression classifier at each time using stratified K-fold CV. We report the area under the receiver operating characteristic curve (AUC-ROC) scores from the out-of-fold probabilities as a measure of the discriminatory power of the model (AUC ≤0.5 indicates random chance; AUC = 1 indicates perfect discrimination). Statistical inference used subject-level label permutation to preserve within-subject temporal dependence: we repeatedly permuted the subject-to-label assignment and, at each time, recomputed AUC and centroid distance to obtain permutation distributions. Two-sided *p*-values were estimated, and Benjamini-Hochberg FDR was applied independently across time for each metric.

## Results

To understand drivers’ behavior and the low-dimensional dynamics underlying autonomous and manual driving, we performed time-wise principal component analysis (PCA) on high-dimensional behavioral data collected during a virtual drive. The analysis was performed on consecutive time segments (subjects × features) on 284 participants described by seven behavioral variables. Given the temporal nature of the data, we analyzed and compared several aspects of the PCA results. We first examined the temporal profile of eigenvalues and changes in explained variance. We then projected the behavioral variables onto the first two principal components (PCs) and assessed each variable’s representation within each PC throughout the drive. We concluded by calculating the data’s effective dimensionality (ED) from its eigenvalue spectrum using various ED estimators, and trained a logistic regression classifier to evaluate the separability of driving conditions using the scores for the first two PCs. This comprehensive analysis of the principal behavioral components extracted via PCA provides valuable insights into key aspects of driver behavior, facilitating comparisons of behavioral patterns between driving conditions.

### Temporal shifts in explained variance across principal components

To understand the underlying structure of driver behavior, we applied PCA on all behavioral variables to reduce the data’s complexity into a set of principal components (PCs). We examined the explained variance associated with each behavioral PC over time to determine the dimensionality and dominant modes of variability in the data. The results show temporal shifts in explained variance across PCs: the first two PCs consistently captured a substantial proportion of the cumulative variance throughout the entire drive (PC1 *M* = 0.273, *SD* = 0.032; and PC2 *M* = 0.187, *SD* = 0.018), often exceeding 50% of the cumulative variance (Fig 4). This pattern was stable both inside and outside critical driving events, indicating that the data can be effectively represented by a low-dimensional structure characterized by the dominant influence of the PCs with the largest eigenvalues. This low-dimensional representation indicates that a large proportion of behavioral variability can be captured by the early components, thereby reducing the complexity inherent in multivariate behavioral data. This suggests that driver behavior is surprisingly low-dimensional, with a few dominant modes of variability governing core behavioral patterns.

**Fig 4.**
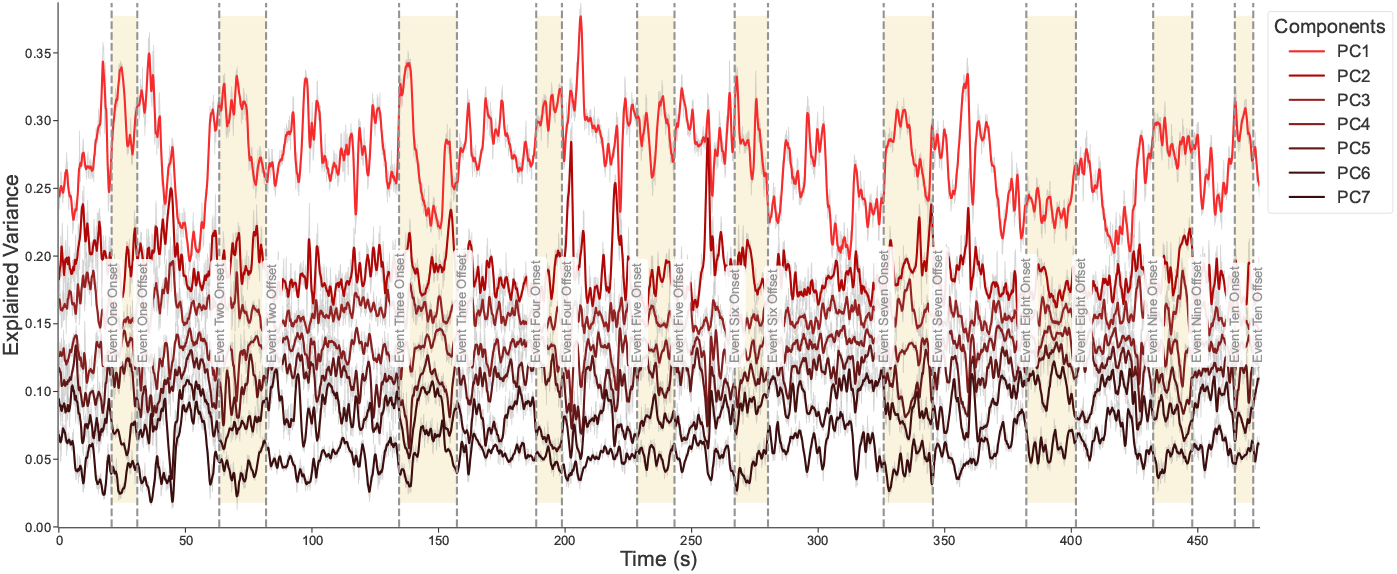
Temporal dynamics of the principal components’ explained variance over the duration of the drive. The thin gray lines denote the actual variance obtained from the dataset, serving as a reference for signal variability. Solid red lines depict the smoothed variance, calculated using a 50-sample rolling window (≈ 1 second), emphasizing trends in variance over time. Yellow windows represent the ten critical events, and dotted gray lines indicate their respective onsets and offsets, illustrating key moments that influence behavioral variance.

Moreover, an increase in the proportion of variance accounted for by the first two PCs during critical events was observed. Across all ten events, PC1 (*M* = 0.319, *SD* = 0.021) and PC2 (*M* = 0.208, *SD* = 0.010) jointly reached, on average, a maximum explained variance exceeding 50% of the cumulative variance. Because a larger share of total variance was thus concentrated in the first two dimensions, the data effectively occupied fewer dimensions during these periods. This concentration of variance in the leading PCs suggests a trend toward more pronounced, low-dimensional behavioral patterns around critical events, reflecting heightened behavioral consistency and stronger engagement of primary driving-related behaviors during risk-related situations. Consequently, behavior is not only low-dimensional overall, but its dimensionality further reduces in context-specific situations.

### Principal component variance reallocates around critical event onset

To quantify how drivers’ multivariate behavior reorganizes around critical events, we compared the distribution of variance before (BO) and after (AO) the onset of each of the ten critical events. Events were aligned to onset, and statistical analysis was performed within a ± 5-second window. The PCs’ variance distributions across all events show a clear variance redistribution (Fig 5). After onset, higher variance consistently concentrates in the early PCs, indicating event-evoked reorganization. Specifically, PC1 captures a larger proportion of variance (AO mean shifts upwards), whereas later PCs capture less (PC6-7 AO mean shifts downwards). Moreover, we found a significant difference between the BO and AO time windows (*χ*^2^(70, *N* = 70) = 23051.27, *p <* 0.001). The pooled changes in variance across all event × PC pairs were small 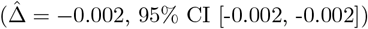. Since the total variance across BO and AO windows is 1, the variance difference indicates an increase in PC1 variance that was compensated by decreases in other PCs. Cochran’s Q redistribution test confirmed that these changes were not uniform across PCs but reflected a reallocation of variance toward the leading component(s) (*Q*(6, *P* = 7) = 7770.18, *p <* 0.001; *I*^2^ = 99.9%). Together, these results indicate a consistent, event-evoked reallocation of variance toward earlier PCs across events, consistent with stronger cross-variable coordination in a lower-dimensional space during hazard response.

**Fig 5.**
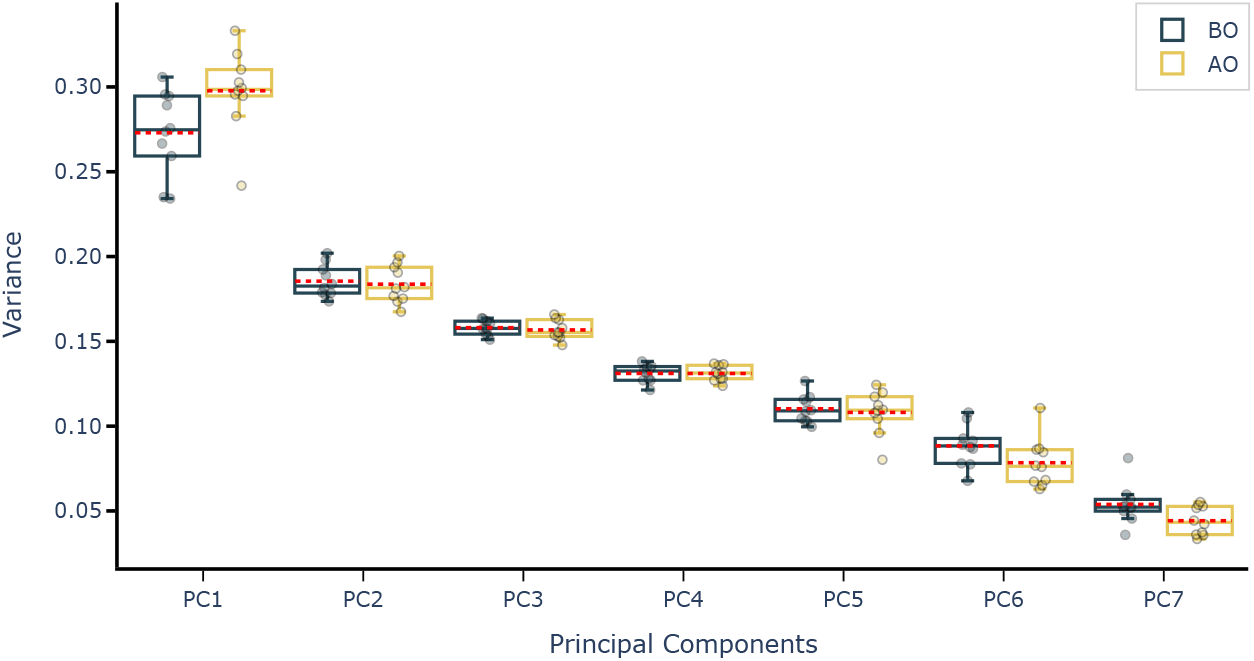
Distribution of variance around critical event onset for all ten events. For each PC, boxes represent the explained variance before (BO) and after (AO) critical event onset. All boxes show the interquartile range, the solid line is the median, the red dotted line marks the mean, and the circles denote the mean variance for each critical event. Higher variance concentrates in early PCs and reallocates after event onset.

### Tracing strategic timepoints of low and high dimensionality based on variance

So far, we have described broad and consistent patterns in the variance and dimensionality of driver behavior across multiple events. However, how behavioral variables relate to each other in the low-dimensional subspace remains unclear. To address this, we need to examine how variables project onto the first two PCs and how their contributions evolve over time. Because the full drive is a long time series, we narrow the focus to a single event to explore the temporal evolution of driving strategies. This allows a detailed examination of how variables interact over time, the mechanisms underlying shifts in dimensionality, and distinct behavioral adaptations that may be obscured when considering aggregated events.

Managing the uncertainty inherent in most traffic situations is challenging, yet drivers often navigate it successfully even in complex scenarios. Understanding how behavior is structured during such critical events is therefore crucial. For this analysis, we focus on an event in which participants confronted a hazard that entered the roadway (S2 Video). Time-wise PCA on eye, head, car, and steering movements revealed a dynamic reallocation of variance over the course of the event (Fig 6). The variance of PC1, for instance, exhibits a clear temporal modulation, starting with a low initial magnitude at onset, followed by a sustained increase to maximum peak, and a subsequent decline (Fig 6A). We traced these strategic timepoints and found that from an initial 45% of cumulative variance explained at onset by the first two PCs (PC1 *eigenvalue* = 1.876, *SD* = 1.369, *variance* = 0.267; PC2 *eigenvalue* = 1.350, *SD* = 1.161, *variance* = 0.192), the system reached a moment of low effective dimensionality between 23.74 and 24.0 seconds. In that window, PC1 (*eigenvalue* = 2.384, *SD* = 1.544, *variance* = 0.339) and PC2 (*eigenvalue* = 1.393, *SD* = 1.180, *variance* = 0.198) jointly accounted for 54% of the cumulative variance. This increase in variance in the early dimensions indicates the emergence of a more structured, low-dimensional behavioral pattern that coincided with clearer separation between autonomous and manual groups (Fig 6B). A subsequent drop in explained variance at around second 28.5 indicated a moment of high effective dimensionality (PC1 *eigenvalue* = 1.884, *SD* = 1.372, *variance* = 0.268; PC2 *eigenvalue* = 1.284, *SD* = 1.133, *variance* = 0.182), suggesting that the system became less structured, variables less coupled, so that more dimensions are required to explain behavior. This pattern is consistent with real-life driving, in which adaptable behavior becomes more or less structured and (condition-) specific through continuous transformations, despite the many DoF involved. These results show that complex behavior can be reduced to a low-dimensional space to identify moments of low and high effective dimensionality, illustrating the adaptive, time-varying structure of behavior around critical events.

**Fig 6.**
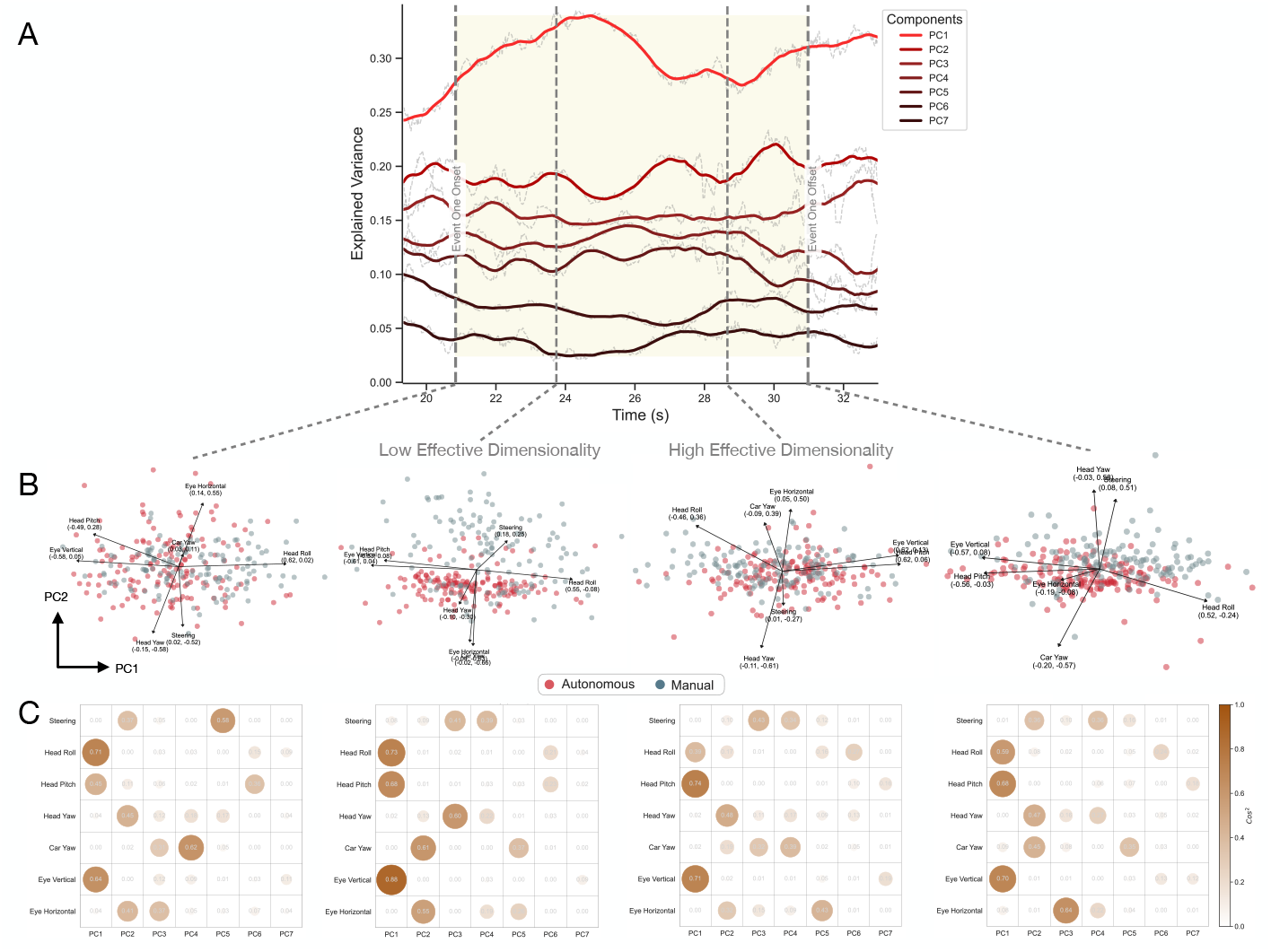
Principal component analysis (PCA) over the course of a critical event. (A) Temporal profile of eigenvalues for PC1 to PC7 before, during, and after a critical event. Dotted gray lines denote the variance for each PC, and solid lines depict the smoothed rolling mean (50-sample window). Vertical gray lines indicate critical event onset, offset, and the times of low and high effective dimensionality. (B) Biplot projections of behavioral variables onto the first two PCs at critical event onset (left), low effective dimensionality (middle-left), high effective dimensionality (middle-right), and critical event offset (right). Each point in the biplot represents a participant belonging to either a manual (blue) or an autonomous (red) driving condition. (C) Square cosine (cos^2^) depicting the quality of representation of all variables on the factor map. Each circle represents the quality of representation of each variable in the corresponding dimension. A high cos^2^ (dark brown) indicates good representation of the variable on the dimension. For any given variable, the sum of the cos^2^ across all dimensions is 1.

### Insights into sensorimotor coordination strategies via biplots

We investigated the structure of these spaces of low and high effective dimensionality by projecting the behavioral variables onto PC1 and PC2 to visualize driver behavior in two dimensions (Fig 6B). This projection shows the relationship between the PCA scores and their loadings, providing a clear representation of how different behaviors couple and revealing correlations between the variables and the first two PCs (see S3 Video). Projecting the data onto the strategic timepoints revealed different coordination strategies that unfold over time. At event onset, the biplot shows that manual and autonomous conditions occupy a shared region of the subspace. PC1 is primarily associated with a positive correlation with head roll (*r* = 0.843, *p <* 0.001) and a negative correlation with eye vertical (*r* = −0.799, *p <* 0.001) and head pitch (*r* = −0.667, *p <* 0.001), indicating that postural stability and vertical gaze control are the dominant factors in this dimension as lateral head tilt co-varies with opposite changes in vertical gaze and head movements. Minor but significant associations were also observed for eye horizontal (*r* = 0.191, *p <* 0.001) and head yaw (*r* = − 0.203, *p <* 0.001). In contrast, PC2 is most strongly correlated with head yaw (*r* = − 0.670, *p <* 0.001) and steering (*r* = − 0.604, *p <* 0.001), establishing this principal behavior as an active steering orientation. This orientation is further defined by a strong, simultaneous opposing horizontal movement of eyes (*r* = 0.639, *p <* 0.001), head pitch (*r* = 0.327, *p <* 0.001), and to a lesser extent car yaw (*r* = 0.127, *p <* 0.05). At event onset, PC2 therefore captures a complex head-centric *“orient-and-scan”* behavior (head turning leads steering): drivers turn their heads and steer in one direction while their gaze scans in the opposite direction, likely to assess alternative paths or monitor the environment. The absence of clear condition separation in either dimension during event onset suggests a common baseline behavior.

These effects change over the course of the event. We therefore examined the timepoint after onset at which the cumulative variance explained by PC1 and PC2 peaked (*low effective dimensionality*; cumulative variance = 54%) (Fig 6A). At this timepoint, manual and autonomous conditions form two clusters with partial overlap, highlighting both shared behavioral (PC1) and condition-specific (PC2) variability in the underlying dynamics (Fig 6B *– low effective dimensionality*). The analysis reveals a significant evolution in the shared behavioral strategy represented by PC1. At event onset, PC1 was primarily characterized by a compensatory relationship between head roll and vertical eye/head movements. As the event progresses, this pattern both strengthens and expands. The underlying coupling represented by PC1 intensifies considerably: the negative correlations of eye vertical (*r* = − 0.937, *p <* 0.001) and head pitch (*r* = − 0.822, *p <* 0.001) with PC1 become much stronger than they were at onset and still co-vary with a comparably strong correlation with head roll (*r* = 0.852, *p <* 0.001). Compared to event onset, eye vertical and head pitch correlate more strongly with PC1 and with each other as indicated by the smaller angles between their vectors (i.e., they co-vary together), and both movement types are negatively correlated with head roll. Notably, the behavioral strategy represented by PC1 expands to incorporate other motor-related variables. It now shows modest but significant correlations with steering (*r* = 0.274, *p <* 0.001) and head yaw (*r* = − 0.154, *p <* 0.05), indicating the emergence of a modest coupling that was absent at onset. At low effective dimensionality, steering and head yaw thus become more temporally coordinated with the shared behavioral dynamics captured by PC1. Importantly, this coupling reflects changes in the co-variation between these variables rather than an increase in the magnitude of their movements, as variables are normalized. This observed effect likely reflects a shift in how steering adjustments and head yaw movements are synchronized within this principal behavioral pattern, rather than an increase in the amplitude of their individual responses.

These after-onset coordination effects on how steering covaries with other behavioral variables become more evident in PC2 at the timepoint of low effective dimensionality. The behavioral dynamics captured by PC2 fundamentally reconfigured into a “coordinated turn execution” mode that separates the driving groups. In a substantial shift from the behavior at onset, PC2 became dominated by a unified coupling of gaze and vehicle heading, showing strong negative correlation with car yaw (*r* = − 0.779, *p <* 0.001) and eye horizontal (*r* = − 0.741, *p <* 0.001). The eyes now move in the direction of the turn rather than against it, indicating a pattern in which drivers look smoothly into the turn as the vehicle turns. Concurrently, the contribution of head yaw decreased after onset (*r* = − 0.354, *p <* 0.001), and now steering became a contrasting, positive correlate (*r* = 0.299, *p <* 0.001), reflecting a change in its covariation with the other variables. In contrast to the head-turning behavior seen at onset, heading now appears to rely less on turning the head and more on vehicle plus eye horizontal movements (vehicle leads). This strengthened coupling between car yaw, eye horizontal, and head yaw suggests that after onset, PC2 becomes an orientation-heading mode driven largely by horizontal movements, with steering acting as a secondary control variable that is particularly prominent for drivers in the manual condition.

This new dynamic, representing active, gaze-coupled control of the vehicle heading, is a characteristic signature of the manual driving condition, causing the driving groups to separate into distinct clusters along the PC2 axis (Fig 6B). Specifically, the biplot at the timepoint of low effective dimensionality shows that, in the normalized PC1–PC2 space, manual drivers differ primarily in how steering coordinates with the other variables, whereas autonomous drivers differ primarily in how horizontal movements coordinate with the rest. The partial overlap between the conditions suggests that, although certain variables contribute to condition separation, other factors (not captured by the included variables) lead to similarities between the conditions. Finally, at high effective dimensionality and at critical event offset, condition separability decreases, similarity scores spread across more dimensions, and behavioral effects become less consistent (see Fig 6B-C and S1 Table). By jointly analyzing the variables’ contributions and their correlations with the first two PCs, we found a gradual emergence of after-onset steering covariation with other variables in PC1 that is expressed more prominently in PC2, leading to group separability. PC2 itself transitions from a head-centric orienting component at onset to a control-related vehicle/gaze-centric component after onset, indicating divergent strategies for vehicle heading control following event onset.

### Variables’ contributions redistribute towards early dimensions after onset

Given the observed shift in variance after onset and the gradual reorganization of variable contributions toward an emerging low-dimensional structure, we assessed whether variables became more strongly represented in the early dimensions. We calculated cosine similarity (cos^2^) to identify which variables contributed most to each dimension and to quantify the quality of their representation (Fig 6C and S3 Video). The similarity matrix indicates a clear pattern of consolidation. In the first dimension, we observed increased similarity for vertical head and eye movements from critical event onset (vertical eye cos^2^ = 0.64, head pitch cos^2^ = 0.45, head roll cos^2^ = 0.71) to the timepoint of low effective dimensionality (vertical eye cos^2^ = 0.88, head pitch cos^2^ = 0.68, head roll cos^2^ = 0.73). Notably, the contributions of vertical eye and head movements indicated a substantial and parallel growth, increasing by +0.24 and +0.23, respectively. This occurred alongside the already-dominant contribution of head roll, which served as a stable anchor for this dimension. The results also showed a modest similarity for steering in the first dimension that was absent at onset (steering cos^2^ = 0.08), consistent with the modest increase in correlation between this dimension and steering reported above. The implication of this coordinated convergence is the formation of a highly efficient eye-head-posture coupling, in which eye and head inputs merge to serve a single, prioritized function during critical events: maintaining vertical orientation.

A more dramatic transformation occurred in the second dimension. At event onset, PC2 was primarily defined by head-centric turning behavior (head yaw cos^2^ = 0.45, steering cos^2^ = 0.37). However, by the timepoint of low effective dimensionality, its contribution pattern had completely shifted. At this timepoint, the similarity of head yaw (cos^2^ = 0.13) and steering (cos^2^ = 0.09) with PC2 decreased drastically by − 0.32 and − 0.28, respectively. Meanwhile, horizontal movements of the car and eyes became dominant. Car yaw increased from cos^2^ = 0.02 at onset to cos^2^ = 0.61, and eye horizontal movements increased from cos^2^ = 0.41 to cos^2^ = 0.55, indicating a dynamic coupling of gaze to the vehicle’s motion. This contrast confirms a shift in the second dimension from a head-centric turning behavior to a vehicle-plus-eyes leading approach. This shift constitutes a functional handover, in which the underlying behavioral dynamics transition from representing the initiation of a turn to representing the execution and monitoring of the vehicle’s trajectory. Similar transformations can be observed at later stages of the event, such as subsequent timepoints of high effective dimensionality and at event offset, where variables’ similarity scores spread across more dimensions (Fig 6C), indicating that more dimensions are needed to achieve a comprehensive representation of the underlying behavior (see S3 Video for a complete reconstruction of the similarity matrix over the course of the event). Crucially, the fact that variables’ similarity consolidates toward early dimensions precisely at the moment of low effective dimensionality demonstrates a temporary compression of the behavior into a more structured, parsimonious state.

### Effective dimensionality declines after critical event onset

Building on the single-event evidence showing that variables’ contributions concentrate in early dimensions after critical event onset, we next sought to quantify the overall simplification of behavior around critical events and to determine whether this simplification generalizes across all event onsets. We hypothesized that the overall dimensionality of the data would be low and would be even lower after event onset. Because the raw eigenspectrum is a complex distribution of many eigenvalues rather than a single measure, we used established estimators to summarize the eigenspectrum into a single, interpretable index that reflects the data’s effective dimensionality (ED). ED quantifies how variance is distributed across dimensions and, consequently, the number of dimensions that are effectively used. We quantified ED using four estimators (*n*_1_, *n*_2_, *n*_∞_, and *n*_*C*_) within a ± 5-second window around each critical event onset, excluding the onset sample itself from statistical analysis. Conceptually, these estimators quantify how “spread out” the variance is: if variance is concentrated in just a few dimensions, ED is low; if it is distributed across many dimensions, ED is high. The subscript for each estimator weights the contribution of dimensions differently: *n*_1_ provides a balanced assessment across all meaningful dimensions, *n*_2_ places greater weight on the dominant dimensions while discounting smaller ones, *n*_∞_ is the extreme case considering only the single largest dimension, and *n*_*C*_ derives dimensionality by subtracting the variance of the eigenvalues from the total number of variables (see Methods section). Here, we report the results for all estimators and focus on *n*_2_ as it effectively discounts the large number of small eigenvalues often associated with increased sensor counts. This robustness makes *n*_2_ well suited for capturing the consolidation of behavior into a structured state without being overly influenced by noise.

To illustrate the effect of a critical event on dimensionality, we first examined the *n*_2_ trajectory for a single representative event. Before the event onset, ED was relatively stable, reaching a maximum of 6.2 dimensions. A sharp and rapid decline in ED began shortly before onset, reaching a minimum of 4.4 dimensions after onset (Fig 7A); a drop of almost 2 effective dimensions. We therefore asked whether this behavioral simplification is systematic across all critical events. The results revealed a clear and consistent pattern of sustained decrease in ED after onset across events (Fig 7B). While individual event trajectories of ED show natural variability, they almost consistently drop below their before-onset baseline. The mean trajectory (black line) confirms a clear tendency for ED to decline immediately after onset, moving from a before-onset average of about 5.8 dimensions to a sustained lower ED after onset. This trend is validated by strong statistical evidence. Event-level chi-squared tests revealed a significant reduction in *n*_2_ between before (BO) and after onset (AO) windows *χ*^2^(10, *N* = 10) = 8025.01, *p <* 0.001, with an inverse-variance-weighted mean difference of 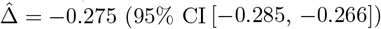, indicating consistent lower ED after onset across events. Furthermore, this event-evoked reduction in ED after onset was consistent across all four metrics (Holm-adjusted *p <* 0.001). Comparable effects were found for *n*_1_, *χ*^2^(10, *N* = 10) = 6269.60, *p <* 0.001, 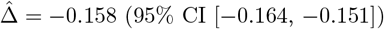, and *n*_∞_ *χ*^2^(10, *N* = 10) = 9493.14, *p <* 0.001,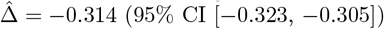. For *n*_*C*_, the effect was smaller but still moderate, 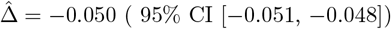, and the global BO and AO difference remained highly significant, *χ*^2^(10, *N* = 10) = 7281.28, *p <* 0.001. These results demonstrate that the simplification of behavior observed in a single event generalizes across events as a robust, stereotyped response in which the system consistently reduces complexity to navigate critical situations.

**Fig 7.**
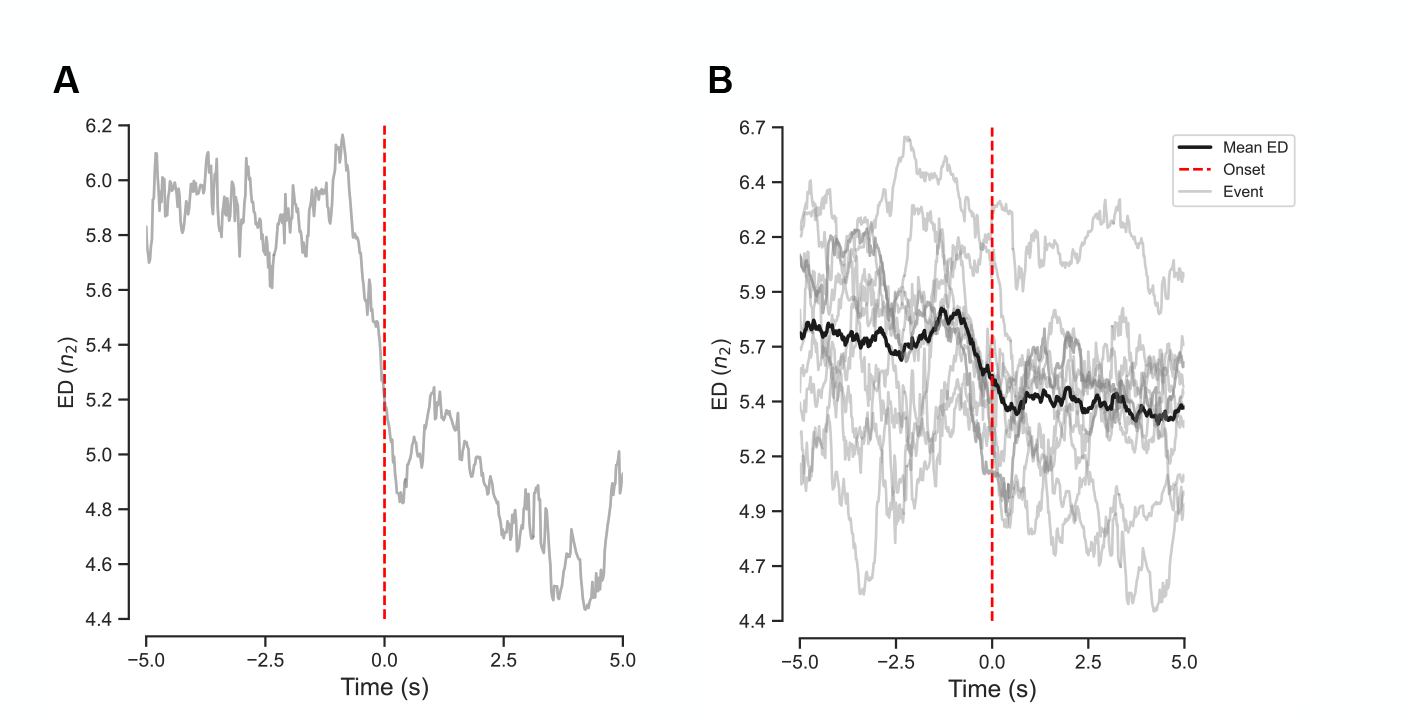
Critical events induce a lower-dimensional behavioral state. (A) The *n*_2_ effective dimensionality trajectory for a single, representative event within a ± 5-second window around onset. The ED remains stable before onset (*t* = 0, red dashed line), then exhibits a sharp and rapid drop, illustrating a clear simplification of behavior. (B) This simplification proves systematic after aligning the ED trajectories for all ten critical events. Individual event trajectories (light gray lines) show natural variability but a consistent tendency to decrease after onset. The mean trajectory (thick black line) reveals a significant and sustained reduction in ED, a trend confirmed by an event-level chi-square test across several ED metrics comparing the before and after onset periods (Holm-adjusted *p <* 0.001).

### Driving conditions have distinct low-dimensional representations within PC1-PC2

After having established the low-dimensional nature of behavior, we next tested whether this simplified PC1-PC2 subspace of multi-signal behavior was functionally meaningful. Specifically, we hypothesized that it would capture distinct behavioral signatures of manual versus autonomous driving, allowing for separation between the two conditions. To achieve this, we assessed both the geometric separability of the conditions and their predictive power using a time-resolved discriminant and classification approach. First, we examined the geometric distance between the two conditions within the PC1-PC2 space. For a representative event, the discriminant scores analysis shows consistent separation between manual (*M* = 0.43, *SD* = 0.14) and autonomous groups (*M* = − 0.43, *SD* = 0.14) with time-sensitive windows where separability increases early after critical event onset (Fig 8A). The centroid distance between the groups confirmed this separation, averaging a centroid distance of 0.86 (*SD* = 0.28; 95% CI: [0.58, 1.15]), indicating a statistically robust separation across time that strengthens during certain periods (FDR ≤0.05) (Fig 8B). This effect was sustained across events with an average centroid distance of 0.80 (*SD* = 0.21; 95% CI: [0.69, 0.93]) (see Table 1). This geometric analysis provides evidence that the low-dimensional representations of manual and autonomous driving conditions captured by the primary PC1-PC2 behavioral subspace correspond to robustly separable states.

**Table 1.** Across-event summary (mean ± SD, 95% bootstrap CI across events).

| Metric | Mean | SD | CI<sub>low</sub> | CI<sub>high</sub> | $n_{\text{events}}$ |
| --- | --- | --- | --- | --- | --- |
| Manual disc. score | 0.40 | 0.10 | 0.34 | 0.46 | 10 |
| Autonomous disc. score | -0.40 | 0.10 | -0.46 | -0.34 | 10 |
| \|Manual – Autonomous\| disc. difference | 0.80 | 0.21 | 0.69 | 0.93 | 10 |
| Centroid distance | 0.80 | 0.21 | 0.69 | 0.93 | 10 |
| AUC at per-event max | 0.87 | 0.07 | 0.83 | 0.91 | 10 |
| Precision@0.5 at max AUC | 0.82 | 0.08 | 0.77 | 0.87 | 10 |
| Recall@0.5 at max AUC | 0.77 | 0.10 | 0.71 | 0.83 | 10 |
| F1@0.5 at max AUC | 0.80 | 0.09 | 0.74 | 0.85 | 10 |
| Max F1 at max AUC | 0.82 | 0.07 | 0.78 | 0.87 | 10 |
| AUC (event means) | 0.67 | 0.05 | 0.65 | 0.71 | 10 |
| Precision@0.5 → event means | 0.64 | 0.05 | 0.61 | 0.67 | 10 |
| Recall@0.5 → event means | 0.52 | 0.08 | 0.47 | 0.56 | 10 |
| F1@0.5 → event means | 0.57 | 0.07 | 0.53 | 0.61 | 10 |
| Max F1 → event means | 0.68 | 0.03 | 0.66 | 0.70 | 10 |

**Fig 8.**
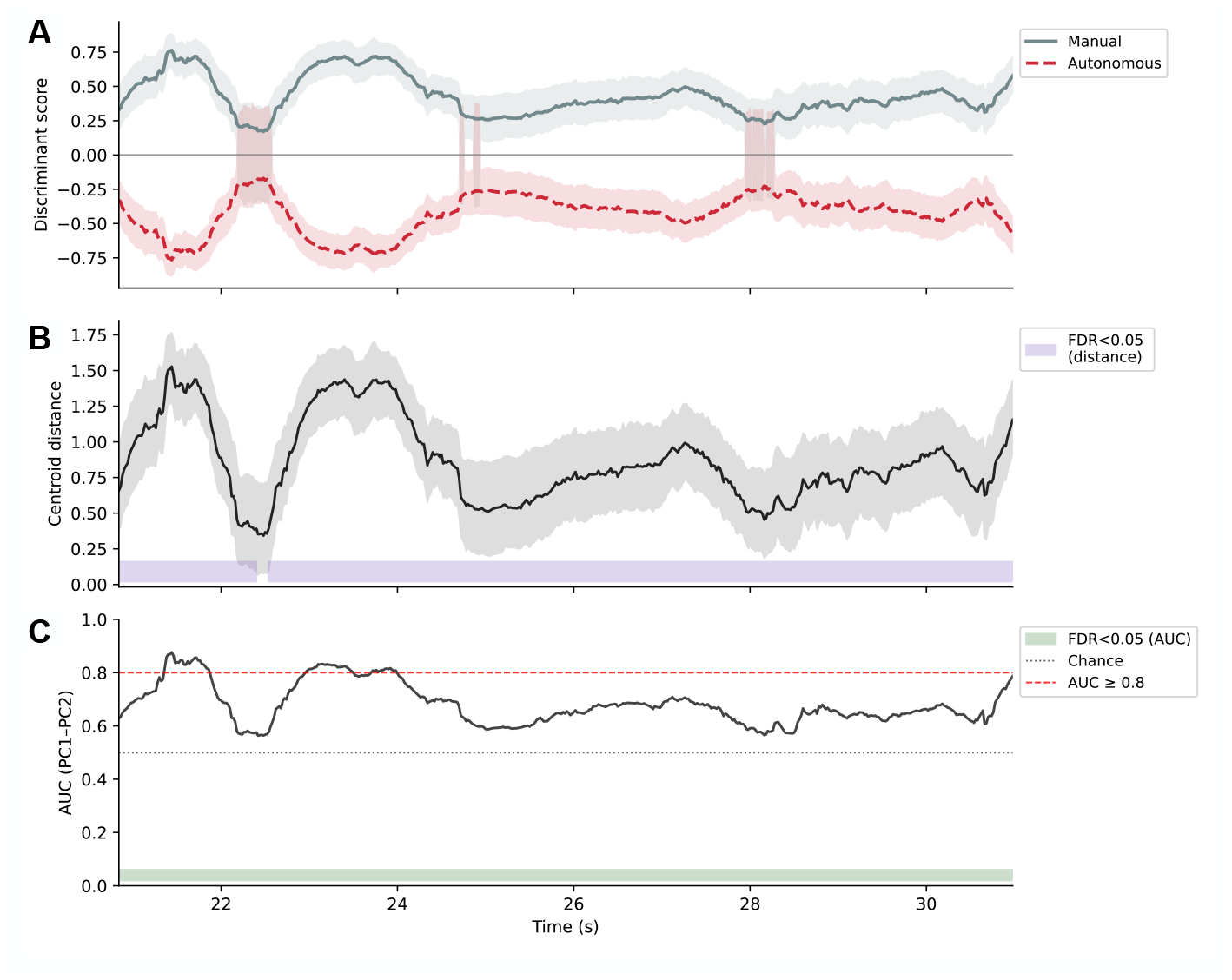
Driving conditions have distinct low-dimensional representations in PC1-PC2 subspace. Three panels showing that the first two principal components (PC1-PC2) of behavior are sufficient to robustly separate manual and autonomous driving conditions. (A) Discriminant scores over time for a representative event show a clear separation between the manual (solid blue) and autonomous (dashed red) conditions. Shaded areas represent 95% bootstrap CIs (resampling participants within each group at each time t), indicating minimal overlap between the two conditions. (B) The geometric distance between the two group centroids (*D* = ∥*c*_Manual_ − *c*_Auto_∥) in the PC1-PC2 space over time with 95% bootstrap CIs (class-stratified resampling at fixed t). The purple bar at the bottom highlights timepoints where this distance is statistically significant (FDR-corrected), as obtained from subject-level label permutations. (C) Predictive power of the PC1-PC2 subspace, measured by the Area Under the Curve (AUC) for a stratified K-fold logistic regression classifier with obtained out-of-fold probabilities. The classifier consistently exceeds chance performance (dotted gray line, AUC = 0.5), with several timepoints achieving high accuracy (dashed red line, AUC ≥ 0.8). The green bar indicates the same subject-level permutation framework. Together, larger discriminant scores and centroid distance co-occur with elevated AUC, indicating transient windows of strong, statistically robust condition separability.

Second, we assessed whether the PC1-PC2 subspace retains enough condition-specific information for a classifier to predict the driving condition. For the same representative event, the classifier performance reached a maximum AUC of 0.87 (precision@0.5 = 0.80, recall@0.5 = 0.77), demonstrating the existence of transient windows of strong separability (Fig 8C). While the average performance over the entire event was lower (AUC = 0.68, *SD* = 0.07), the classifier’s discriminatory power remained significantly above chance (FDR ≤0.05). Across all events (*N* = 10), the classifier consistently performed above chance with an average AUC of 0.67 (*SD* = 0.05; 95% CI: [0.64, 0.70]). Crucially, when examining the timepoint of maximum discriminability within each event, the classifier achieved a mean peak AUC of 0.87 (*SD* = 0.07; 95% CI: [0.82, 0.91]). This indicates that, while the exact timing may vary, each critical event contains a clear, transient window in which the behavioral signatures of the two driving conditions are highly distinct in the PC1-PC2 subspace. Together, these geometric and predictive analyses demonstrate that major behavioral differences between manual and autonomous driving are captured within just the first two PCs.

These findings support the idea that the low-dimensional subspace is not merely a statistical summary but a functionally relevant representation of driver behavior.

## Discussion

This study aimed to examine the low-dimensional dynamics of complex, natural behavior. Naturalistic driving in virtual reality revealed that a small set of latent dimensions captures most of the variance in eye, head, and vehicle behavior, and that this effective dimensionality reduces further around hazardous events. Throughout the drive, the first two principal components (*eigenbehaviors*) consistently captured most of the variance, frequently exceeding half of the total variance. Established estimators of effective dimensionality further supported the idea that only a few dimensions were effectively used [24, 34]. Around critical events, variance reallocates toward the leading components, and effective dimensionality drops. Consequently, the contributions of the behavioral variables reorganized: early after onset, vertical eye movements and head pitch dominated alongside head roll, whereas slightly later, horizontal eye and vehicle heading became more pronounced and the coordination between steering and the other behavioral variables became a defining feature of the low-dimensional structure, particularly for manual drivers. Within this compressed subspace, manual and autonomous driving groups occupied overlapping but reliably separable regions. Discriminant and logistic regression classification analyses based on the first two principal components supported these findings, demonstrating that both driving conditions occupied clear, separate regions in the PC1-PC2 subspace, with significantly above-chance classification accuracy and strong separability during transient time windows. Together, these findings indicate that the visuomotor coordination underlying driving behavior is both low-dimensional and context-sensitive, with hazard-related compression revealing group-specific coordination patterns.

These results provide a naturalistic, task-level perspective on the long-standing degrees-of-freedom problem [1, 3, 6]. Previous work on motor synergies has shown that coordinated action in grasping and locomotion can be described in terms of low-dimensional modules [39, 43–47], and that neural population activity evolves on low-dimensional manifolds supporting movement generation [29, 36, 37, 49]. In line with these frameworks, treating behavior as trajectories on a structured manifold, the observed reduction of effective dimensionality around critical events suggests that the system temporarily collapses onto a smaller set of coordinated modes that channel visuomotor variability into task-relevant degrees of freedom. This extends dimensionality reduction approaches that have been predominantly applied to animal behavior and repetitive human tasks [18–23, 52] to complex, multi-effector human behavior. Taken together, these findings indicate that the driver-vehicle system forms a functional unit [65] whose behavior can be understood in terms of a small set of context-sensitive coordination patterns.

Analyzing such high-dimensional data requires specific choices regarding how the multidimensional data structure (e.g., participants × features × time) is decomposed, as the matricization strategy dictates the biological interpretation of the resulting manifold. Here, we applied a time-resolved, cross-sectional strategy: at each timepoint we stacked participants to form a subjects × features matrix. Consequently, the PCA extracts directions of maximal inter-individual covariation, yielding a group-level manifold that captures shared visuomotor coordination strategies and enables direct comparison of behavioral modes across individuals [15]. This contrasts with approaches that leverage repeated trial structure, such as that of Keshava et al. [54]. In their work involving 4-dimensional data (features × time × participants), PCA was applied across repetitions of similar actions within individual subjects. That approach characterizes intra-individual trial-to-trial variability, highlighting how the eye, head, and hand couple synergistically to support consistent action planning despite motor noise. While such “repetition slicing” summarizes the consistency of motor execution, our cross-sectional approach captures the structural variability of coordination strategies across the population. This distinction is fundamental for interpretation: because our principal components are defined by between-participant variance, separation in the manifold reflects differences in style (e.g., how steering covaries with gaze and head orientation) rather than trial-by-trial fluctuations in execution. Thus, our approach reveals how different drivers solve the same motor problem, rather than how a single driver stabilizes a movement across attempts.

Our results support the central hypothesis that visuomotor coordination during driving is constrained to a low-dimensional manifold and that critical events further compress behavior into an even lower-dimensional subspace. The variance explained by the first two principal components fluctuates throughout the drive, occasionally exceeding 50% of the total variance. This aligns with previous findings indicating that manifolds for unconstrained behaviors exhibit slightly higher dimensionality than those of constrained behavior, with low-dimensional manifolds carrying task-relevant information [26]. The variability of this task-relevant information has been shown to be smaller than that of task-irrelevant information [66]. Consistent with a shift toward such a task-relevant, low-dimensional regime, we observed two complementary patterns around critical event onsets: the reorganization of variables’ contributions toward the early dimensions and a consistent, significant decrease in effective dimensionality across all events. Furthermore, the obtained subspace carries condition-specific information that differentiates between driving groups. Manual driving was characterized by steering-dominant coordination patterns within the low-dimensional space. In contrast, autonomous driving was defined by the covariance structure of horizontal gaze and car heading, consistent with drivers monitoring the environment and the automated system rather than controlling the vehicle. This finding aligns with previous research demonstrating that the underlying visuomotor coordination for steering persists even when vehicle control is automated [67]. The discriminant analysis and classification performance demonstrate that a small number of latent dimensions is sufficient to capture these differences, suggesting that the obtained manifold reflects meaningful functional structure. This indicates that adaptive, context-dependent reductions in dimensionality are a hallmark of skilled behavior, rather than simply by-products of biomechanical constraints.

The issue of developing methodologies to capture and analyze the structure of complex human behavior in natural settings remains relevant. Despite the large sample and rich behavioral dataset, certain limitations constrain the extent to which our findings can address broader questions regarding the dimensionality of visuomotor behavior. First, although principal component analysis (PCA) has proven useful for uncovering low-dimensional structures across behavioral and neural data [12, 39, 49, 68], the debate persists over whether linear or more complex nonlinear dimensionality reduction techniques are better suited for capturing the underlying dynamics [27, 69]. Indeed, independent time-wise PCA analyses ignore temporal dependencies in the behavioral signals, treating each time slice as statistically independent. This choice allows a straightforward, interpretable characterization of moment-to-moment coordination patterns, but fails to capture how coordination evolves through time and its underlying continuous functional form. To understand these spatial and temporal structures, future work could examine more temporally aware methods, such as functional PCA [70–72] and other nonlinear methods [26, 73–75]. These methods could, for instance, help elucidate how task-relevant components are modulated over time and by task demands.

Second, although a representative set of eye, head, and vehicle variables was used, additional factors might prove valuable in investigating the effective dimensionality of driving behavior. These might include pedal forces, body postures, or variables explicitly related to decision-making. These variables likely contribute additional information and could help explain residual variability and the partial overlap among driving groups. Third, although the study focused on understanding the dimensionality of driving behavior around critical events, additional low-dimensional timepoints also arise outside critical events. Visual inspection suggested that these were mostly driven by external scene constraints, such as driving through a tunnel. This directly relates to the interpretation of effective dimensionality. Since effective dimensionality was computed from the eigenvalue spectrum, it quantifies the shared variance among variables rather than the intrinsic importance of an event. In a tunnel segment, for instance, the environment constrains possible movements and, consequently, behavior is stereotyped (e.g., looking straight ahead, uniform steering). Thus, low effective dimensionality should be interpreted as an indicator of strong coupling between effectors given the current environmental demands, rather than as a direct marker of danger or difficulty. Fully addressing the relationship between effective dimensionality and task demands will require modeling the environment and task structure more explicitly, which is beyond the scope of the present study.

Despite these limitations, this study demonstrates that complex, unconstrained visuomotor coordination during driving is confined to a low-dimensional space whose effective dimensionality dynamically decreases around hazards. It shows that effective dimensionality provides a compact, interpretable metric of how many dimensions behavior effectively occupies and how tightly effectors are coupled over time. This metric seems to decrease for both hazard-related coordination and more mundane episodes of environmental constraints. Moreover, this study establishes that applying PCA across individuals yields a common subspace in which manual and autonomous driving groups leave distinct signatures. This suggests that low-dimensional behavioral dynamics could serve as a bridge between human-centered motor control theories and the design of autonomous systems. Consistent with the view that cognition is inherently situated and temporal [76], our results highlight the importance of modeling not only what drivers do, but also how the structure of their behavior evolves in a task-dependent manner as the environment changes.

## Conclusion

In summary, this study demonstrates that, despite their inherent complexity and variability, natural behaviors such as driving can be captured in a compact, low-dimensional behavioral space. Across nearly three hundred participants and ten hazardous events, behavior occupied a shared, low-dimensional subspace, and its effective dimensionality decreased systematically around hazards. This subspace carries condition-specific information that enables reliable separation of driving modes. The transient reductions in effective dimensionality reflect moments in which perception and action are tightly coupled, with task-relevant coordination patterns dominating idiosyncratic variability. Manual and autonomous driving behavior occupied a common manifold yet traced distinct trajectories within it, suggesting that the delegation of control systematically reshapes visuomotor coordination while preserving overall structure. Finally, investigating the low-dimensional structure of unconstrained behavior will be crucial for comparing human and machine interaction strategies and for designing autonomous systems that are receptive to the context-dependent coordination patterns that characterize human behavior.

## Supporting information

**S1 Video. Exemplary driving events**. A video showing exemplars of various critical events, including the training scene, a lane closure, a roadway obstruction, and a pedestrian intrusion. [S1_video_exemplary_driving_events.mp4](https://osf.io/3y4ea/files/osfstorage/6998c2331dc3e3e6a05061be)

**S2 Video. Four-panel composite of a driving hazard**. Synchronized view of a single driving event. (Top left) exemplar video of the drive. (Top right) cosine scores. (Bottom left) explained variance of each principal component over time (seconds). (Bottom right) PCA biplot. All panels are aligned to a common time axis. A slow-motion effect was applied within the hazard-related time window. The composite illustrates how temporal changes in variance and loadings relate to the driving behavior. [S2_video_four-panel_composite.mp4](https://osf.io/3y4ea/files/osfstorage/6998c2b9b2b00dabf2259fca)

**S3 Video. Biplots and cosine matrix for a driving event**. A video showing: (left) the projections of the behavioral variables onto the first two principal components (PC1 and PC2) over time for an exemplary event; (right) the cosine similarity of the variables with the components. [S3_video_biplots_cosplots_event1.mp4](https://osf.io/3y4ea/files/osfstorage/6998c303b2b00dabf2259fe4)

**S1 Table.**
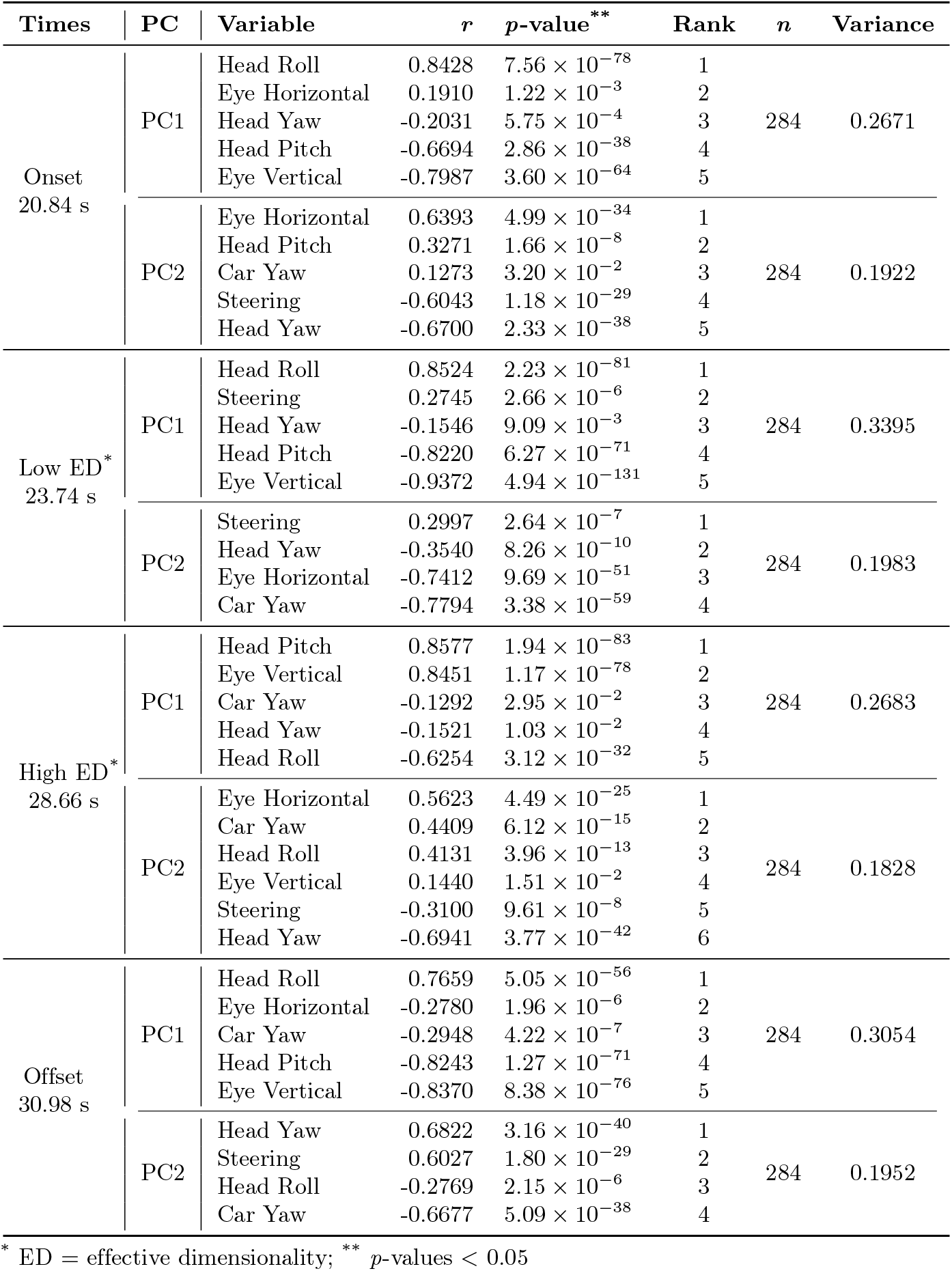
Variables’ correlations with PC1 and PC2. A table containing the correlations of the variables with the first two principal components at different timepoints of the drive during event one; *p*-values, ranks, and cumulative variance are also provided.

## Acknowledgments

We want to thank everyone who contributed to this project within the Unity development team from the Neurobiopsychology and Neuroinformatics departments at the University of Osnabrück. In particular, we thank Farbod Nosrat Nezami, Maximilian Alexander Wächter, and Gordon Pipa for their help in constructing the experimental setup and virtual environment. We also want to thank the German Research Foundation DFG which supported this project in the context of funding the Research Training Group “Situated Cognition” (GRK 274877981).

## Author Contributions

S.D., J.M.-C., and P.K. conceptualized the research; S.D. collected the data; S.D. and J.M.-C. preprocessed and cleaned the data; J.M.-C. analyzed and visualized the data; J.M.-C. wrote the original manuscript draft; J.M.-C. and P.K. reviewed and edited the manuscript. P.K. supervised the project.

## Data Availability

The code for the virtual environment is available GitHub repository under a Creative Commons license https://github.com/Westdrive-Workgroup/LoopAR-public.

Additionally, raw and cleaned data, analysis scripts, and supplementary videos are accessible on the Open Science Framework (OSF) under a Creative Commons Attribution 4.0 International (CC-BY 4.0) at DOI: 10.17605/OSF.IO/B4XKN.

## References

1. Bernstein NA. The Co-ordination and Regulation of Movements. Oxford, UK: Pergamon Press; 1967. Available from: https://api.semanticscholar.org/CorpusID:62634931.

2. Lashley KS. Integrative functions of the cerebral cortex. Physiological Reviews. 1933;13(1):1–42. doi:10.1152/physrev.1933.13.1.1.

3. Profeta VLS, Turvey MT. Bernstein’s levels of movement construction: A contemporary perspective. Human Movement Science. 2018 Feb;57:111–33. doi:10.1016/j.humov.2017.11.013.

4. Kandel ER, Koester JD, Mack SH, Siegelbaum SA. Principles of Neural Science. 6th ed. New York, NY: McGraw Hill Professional; 2021.

5. Turvey MT. Action and perception at the level of synergies. Human Movement Science. 2007 Aug;26(4):657–97. doi:10.1016/j.humov.2007.04.002.

6. Turvey MT, Fonseca S. Nature of Motor Control: Perspectives and Issues. In: Sternad D, editor. Progress in Motor Control: A Multidisciplinary Perspective. Boston, MA: Springer US; 2009. p. 93–123. doi:10.1007/978-0-387-77064-26.

7. Churchland MM, Afshar A, Shenoy KV. A Central Source of Movement Variability. Neuron. 2006 Dec;52(6):1085–96. doi:10.1016/j.neuron.2006.10.034.

8. Osborne LC, Lisberger SG, Bialek W. A sensory source for motor variation. Nature. 2005 Sep;437(7057):412–6. doi:10.1038/nature03961.

9. Fernandes O. Methods for Analyzing Movement Variability. In: Parziale A, Diaz M, Melo F, editors. Graphonomics in Human Body Movement. Bridging Research and Practice from Motor Control to Handwriting Analysis and Recognition. Cham: Springer Nature Switzerland; 2023. p. 191–202. doi:10.1007/978-3-031-45461-514.

10. Frére J, Hug F. Between-subject variability of muscle synergies during a complex motor skill. Frontiers in Computational Neuroscience. 2012;6:99. doi:10.3389/fncom.2012.00099.

11. Peterson SM, Singh SH, Wang NXR, Rao RPN, Brunton BW. Behavioral and Neural Variability of Naturalistic Arm Movements. eNeuro. 2021;8(3):1–15. doi:10.1523/ENEURO.0007-21.2021.

12. Sili D, De Giorgi C, Pizzuti A, Spezialetti M, de Pasquale F, Betti V. The spatio-temporal architecture of everyday manual behavior. Scientific Reports. 2023;13(1):9451. doi:10.1038/s41598-023-36280-4.

13. Stetter BJ, Herzog M, Möhler F, Sell S, Stein T. Modularity in Motor Control: Similarities in Kinematic Synergies Across Varying Locomotion Tasks. Frontiers in Sports and Active Living. 2020;2:596063. doi:10.3389/fspor.2020.596063.

14. Tessari F, West AM, Hogan N. A Geometric Approach for the Comparison of Kinematic Synergy Postures. In: 2025 International Conference on Rehabilitation Robotics (ICORR); 2025. p. 36–43. doi:10.1109/ICORR66766.2025.11063213.

15. Winner TS, Rosenberg MC, Jain K, Kesar TM, Ting LH, Berman GJ. Discovering individual-specific gait signatures from data-driven models of neuromechanical dynamics. PLOS Computational Biology. 2023;19(10):e1011556. doi:10.1371/journal.pcbi.1011556.

16. Berman GJ. Measuring behavior across scales. BMC Biology. 2018;16(1):23. doi:10.1186/s12915-018-0494-7.

17. Gomez-Marin A, Paton JJ, Kampff AR, Costa RM, Mainen ZF. Big behavioral data: psychology, ethology and the foundations of neuroscience. Nature Neuroscience. 2014;17(11):1455–62. doi:10.1038/nn.3812.

18. Berman GJ, Choi DM, Bialek W, Shaevitz JW. Mapping the stereotyped behaviour of freely moving fruit flies. Journal of the Royal Society Interface. 2014;11(99):20140672. doi:10.1098/rsif.2014.0672.

19. Datta SR, Anderson DJ, Branson K, Perona P, Leifer A. Computational Neuroethology: A Call to Action. Neuron. 2019;104(1):11–24. doi:10.1016/j.neuron.2019.09.038.

20. Krakauer JW, Ghazanfar AA, Gomez-Marin A, MacIver MA, Poeppel D. Neuroscience Needs Behavior: Correcting a Reductionist Bias. Neuron. 2017;93(3):480–90. doi:10.1016/j.neuron.2016.12.041.

21. Mathis A, Mamidanna P, Cury KM, Abe T, Murthy VN, Mathis MW, et al. DeepLabCut: markerless pose estimation of user-defined body parts with deep learning. Nature Neuroscience. 2018;21(9):1281–9. doi:10.1038/s41593-018-0209-y.

22. Stephens GJ, Johnson-Kerner B, Bialek W, Ryu WS. Dimensionality and Dynamics in the Behavior of C. elegans. PLOS Computational Biology. 2008;4(4):e1000028. doi:10.1371/journal.pcbi.1000028.

23. Wiltschko AB, Johnson MJ, Iurilli G, Peterson RE, Katon JM, Pashkovski SL, et al. Mapping Sub-Second Structure in Mouse Behavior. Neuron. 2015;88(6):1121–35. doi:10.1016/j.neuron.2015.11.031.

24. Bialek W. On the dimensionality of behavior. Proceedings of the National Academy of Sciences. 2022;119(4):e2021860119. doi:10.1073/pnas.2021860119.

25. Allen PK, Ciocarlie M, Goldfeder C. Grasp Planning Using Low Dimensional Subspaces. In: Balasubramanian R, Santos VJ, editors. The Human Hand as an Inspiration for Robot Hand Development. Cham: Springer; 2014. p. 531–63. doi:10.1007/978-3-319-03017-324.

26. Altan E, Ma X, Miller LE, Perreault EJ, Solla SA. Low-dimensional neural manifolds for the control of constrained and unconstrained movements [Preprint]. bioRxiv; 2023. Available from: https://www.biorxiv.org/content/10.1101/2023.05.25.542264v1.

27. Cui PH, Visell Y. Linear and nonlinear subspace analysis of hand movements during grasping. In: 2014 36th Annual International Conference of the IEEE Engineering in Medicine and Biology Society. IEEE; 2014. p. 2529–32. doi:10.1109/EMBC.2014.6944137.

28. Hachaj T, Ogiela MR, Koptyra K. Human actions modelling and recognition in low-dimensional feature space. In: 2015 10th International Conference on Broadband and Wireless Computing, Communication and Applications (BWCCA). IEEE; 2015. p. 247–54. doi:10.1109/BWCCA.2015.15.

29. Kobak D, Brendel W, Constantinidis C, Feierstein CE, Kepecs A, Mainen ZF, et al. Demixed principal component analysis of neural population data. eLife. 2016;5:e10989. doi:10.7554/eLife.10989.

30. Sanger TD. Human arm movements described by a low-dimensional superposition of principal components. The Journal of Neuroscience. 2000;20(3):1066–72. doi:10.1523/JNEUROSCI.20-03-01066.2000.

31. Abdi H, Williams LJ. Principal component analysis. Wiley interdisciplinary reviews: computational statistics. 2010;2(4):433–59. doi:10.1002/wics.101.

32. Jolliffe IT. Principal Component Analysis. 2nd ed. Springer; 2002. doi:10.1007/b98835.

33. Jolliffe IT, Cadima J. Principal component analysis: a review and recent developments. Philosophical Transactions of the Royal Society A: Mathematical, Physical and Engineering Sciences. 2016;374(2065):20150202. doi:10.1098/rsta.2015.0202.

34. Del Giudice M. Effective dimensionality: A tutorial. Multivariate Behavioral Research. 2021;56(4):527–42. doi:10.1080/00273171.2020.1743631.

35. Dörfler M, Luef F, Skrettingland E. Local structure and effective dimensionality of time series data sets. Applied and Computational Harmonic Analysis. 2024;73:101692. doi:10.1016/j.acha.2024.101692.

36. Leo A, Handjaras G, Bianchi M, Marino H, Gabiccini M, Guidi A, et al. A synergy-based hand control is encoded in human motor cortical areas. eLife. 2016;5:e13420. doi:10.7554/eLife.13420.

37. Pang R, Lansdell BJ, Fairhall AL. Dimensionality reduction in neuroscience. Current Biology. 2016;26(15):R656–60. doi:10.1016/j.cub.2016.05.029.

38. Pellegrino A, Stein H, Cayco-Gajic NA. Dimensionality reduction beyond neural subspaces with slice tensor component analysis. Nature Neuroscience. 2024;27(6):1199–210. doi:10.1038/s41593-024-01626-2.

39. Santello M, Baud-Bovy G, Jörntell H. Neural bases of hand synergies. Frontiers in Computational Neuroscience. 2013;7:23. doi:10.3389/fncom.2013.00023.

40. Kamboj A, Ranganathan R, Tan X, Srivastava V. Human motor learning dynamics in high-dimensional tasks. PLOS Computational Biology. 2024;20(10):e1012455. doi:10.1371/journal.pcbi.1012455.

41. Oshima H, Aoi S, Funato T, Tsujiuchi N, Tsuchiya K. Variant and invariant spatiotemporal structures in kinematic coordination to regulate speed during walking and running. Frontiers in Computational Neuroscience. 2019;13:63. doi:10.3389/fncom.2019.00063.

42. Boe D, Portnova-Fahreeva AA, Sharma A, Rai V, Sie A, Preechayasomboon P, et al. Dimensionality reduction of human gait for prosthetic control. Frontiers in Bioengineering and Biotechnology. 2021;9:724626. doi:10.3389/fbioe.2021.724626.

43. Dominici N, Ivanenko YP, Cappellini G, d’Avella A, Mondí V, Cicchese M, et al. Locomotor primitives in newborn babies and their development. Science. 2011;334(6058):997–9. doi:10.1126/science.1210617.

44. Ivanenko YP, Cappellini G, Dominici N, Poppele RE, Lacquaniti F. Modular control of limb movements during human locomotion. The Journal of Neuroscience. 2007;27(41):11149–61. doi:10.1523/JNEUROSCI.2644-07.2007.

45. d’Avella A, Portone A, Fernandez L, Lacquaniti F. Control of fast-reaching movements by muscle synergy combinations. The Journal of Neuroscience. 2006;26(30):7791–810. doi:10.1523/JNEUROSCI.0830-06.2006.

46. Santello M, Flanders M, Soechting JF. Patterns of Hand Motion during Grasping and the Influence of Sensory Guidance. The Journal of Neuroscience. 2002;22(4):1426–35. doi:10.1523/JNEUROSCI.22-04-01426.2002.

47. Vinjamuri R, Sun M, Chang CC, Lee HN, Sclabassi RJ, Mao ZH. Dimensionality reduction in control and coordination of the human hand. IEEE Transactions on Biomedical Engineering. 2010;57(2):284–95. doi:10.1109/TBME.2009.2032532.

48. Altan E, Solla SA, Miller LE, Perreault EJ. Estimating the dimensionality of the manifold underlying multi-electrode neural recordings. PLOS Computational Biology. 2021;17(10):e1008591. doi:10.1371/journal.pcbi.1008591.

49. Gallego JA, Perich MG, Naufel SN, Ethier C, Solla SA, Miller LE. Cortical population activity within a preserved neural manifold underlies multiple motor behaviors. Nature Communications. 2018;9(1):4233. doi:10.1038/s41467-018-06560-z.

50. Koh N, Ma Z, Sarup A, Kristl AC, Agrios M, Young M, et al. Selective direct influence of motor cortex on limb muscle activity during naturalistic climbing in mice. Nature Neuroscience. 2025;28:2537–49. doi:10.1038/s41593-025-02093-z.

51. Paulus MP, Dulawa SC, Ralph RJ, Geyer MA. Behavioral organization is independent of locomotor activity in 129 and C57 mouse strains. Brain Research. 1999;835(1):27–36. doi:10.1016/S0006-8993(99)01137-3.

52. Marques JC, Lackner S, Félix R, Orger MB. Structure of the Zebrafish locomotor repertoire revealed with unsupervised behavioral clustering. Current Biology. 2018;28(2):181-95.e5. doi:10.1016/j.cub.2017.12.002.

53. Iwaniuk AN, Hurd PL. The evolution of cerebrotypes in birds. Brain, Behavior and Evolution. 2005;65(4):215–30. doi:10.1159/000084313.

54. Keshava A, Wächter MA, Boße F, Schüler T, König P. Low-dimensional representations of visuomotor coordination in natural behavior [Preprint]. bioRxiv; 2024. Available from: 10.1101/2024.03.30.587357.

55. Gibson JJ, Crooks LE. A theoretical field-analysis of automobile-driving. The American Journal of Psychology. 1938;51(3):453–71. doi:10.2307/1416145.

56. Liu H, Taniguchi T, Tanaka Y, Takenaka K, Bando T. Visualization of driving behavior based on hidden feature extraction by using deep learning. IEEE Transactions on Intelligent Transportation Systems. 2017;18(9):2477–89. doi:10.1109/TITS.2017.2649541.

57. Veit F, Heinen T. The role of visual and auditory information in the observation and evaluation of complex skills in Gymnastics. Journal of Physical Fitness, Medicine & Treatment in Sports. 2019;6(2):555683. doi:10.19080/jpfmts.2018.05.555683.

58. Derakhshan S, Nosrat Nezami F, Wächter MA, Stephan A, Pipa G, König P. A Situated Inspection of Autonomous Vehicle Acceptance - A Population Study in Virtual Reality. International Journal of Human-Computer Interaction. 2024:1–20. doi:10.1080/10447318.2024.2358577.

59. Nezami FN, Wächter MA, Maleki N, Spaniol P, Kühne LM, Haas A, et al. Westdrive X LoopAR: an open-access virtual reality project in Unity for evaluating user interaction methods during takeover requests. Sensors. 2021;21(5):1879. doi:10.3390/s21051879.

60. Lê S, Josse J, Husson F. FactoMineR: an R package for multivariate analysis. Journal of Statistical Software. 2008;25(1):1–18. doi:10.18637/jss.v025.i01.

61. Van Lingen HJ, Suarez-Diez M, Saccenti E. Normalization of gene counts affects principal components-based exploratory analysis of RNA-sequencing data. Biochimica et Biophysica Acta (BBA) - Gene Regulatory Mechanisms. 2024;1867(9):195058. doi:10.1016/j.bbagrm.2024.195058.

62. Dam EB, Koch M, Lillholm M. Quaternions, interpolation and animation. vol. 2. Datalogisk Institut, Københavns Universitet Copenhagen, Denmark; 1998. Available from: https://api.semanticscholar.org/CorpusID:115207743.

63. Burul A. Coordinate Systems of 3D Applications Guide [Internet]. San Francisco, CA: Medium; 2022 Feb 14 [cited 26 October 2025]. Medium. Available from: https://ahmetburul.medium.com/coordinate-systems-of-3d-applications-guide-ddfa2194ed88.

64. Unity Technologies. Unity Scripting API: Transform.eulerAngles [Internet]. San Francisco, CA: Unity Technologies; 2024 [cited 26 October 2025]. Unity Documentation Version 6000.0. Available from: https://docs.unity3d.com/6000.0/Documentation/ScriptReference/Transform-eulerAngles.html.

65. Ziv G. An embodied and ecological approach to skill acquisition in racecar driving. Frontiers in Sports and Active Living. 2023;5:1095639. doi:10.3389/fspor.2023.1095639.

66. Furuki D, Takiyama K. Decomposing motion that changes over time into task-relevant and task-irrelevant components in a data-driven manner: application to motor adaptation in whole-body movements. Scientific Reports. 2019;9(1):7246. doi:10.1038/s41598-019-43558-z.

67. Navarro J, Hernout E, Osiurak F, Reynaud E. On the nature of eye-hand coordination in natural steering behavior. PloS one. 2020;15(11):e0242818. doi:10.1371/journal.pone.0242818.

68. Soldado-Magraner J, Mante V, Sahani M. Inferring context-dependent computations through linear approximations of prefrontal cortex dynamics. Frontiers in Bioengineering and Biotechnology. 2024;10(14):eadj0860. doi:10.3389/fbioe.2020.00429.

69. Portnova-Fahreeva AA, Rizzoglio F, Nisky I, Casadio M, Mussa-Ivaldi FA, Rombokas E. Linear and non-linear dimensionality-reduction techniques on full hand kinematics. Frontiers in Bioengineering and Biotechnology. 2020;8:429. doi:10.3389/fbioe.2020.00429.

70. Anirudh R, Turaga P, Su J, Srivastava A. Elastic functional coding of human actions: From vector-fields to latent variables. In: 2015 IEEE Conference on Computer Vision and Pattern Recognition (CVPR); 2015. p. 3147–55. doi:10.1109/CVPR.2015.7298934.

71. Dai W, Sun Y, Qian X. Functional analysis of grasping motion. In: 2013 IEEE/RSJ International Conference on Intelligent Robots and Systems. IEEE; 2013. p. 3507–13. doi:10.1109/IROS.2013.6696856.

72. Kanzler CM, Averta G, Schwarz A, Held JPO, Gassert R, Bicchi A, et al. A low-dimensional representation of arm movements and hand grip forces in post-stroke individuals. Scientific Reports. 2022;12(1):7601. doi:10.1038/s41598-022-11806-4.

73. Furuki D, Takiyama K. A data-driven approach to decompose motion data into task-relevant and task-irrelevant components in categorical outcome. Scientific Reports. 2020;10(1):2422. doi:10.1038/s41598-020-59257-z.

74. Inoue M, Furuki D, Takiyama K. Detecting task-relevant spatiotemporal modules and their relation to motor adaptation. PLOS ONE. 2022;17(10):e0275820. doi:10.1371/journal.pone.0275820.

75. Takiyama K, Yokoyama H, Kaneko N, Nakazawa K. Speed-dependent and mode-dependent modulations of spatiotemporal modules in human locomotion extracted via tensor decomposition. Scientific Reports. 2020;10(1):680. doi:10.1038/s41598-020-57513-w.

76. Vogel DHV, Jording M, Kupke C, Vogeley K. The Temporality of Situated Cognition. Frontiers in Psychology. 2020;11:546212. doi:10.3389/fpsyg.2020.546212.

